# Simultaneous orientation and 3D localization microscopy with a Vortex point spread function

**DOI:** 10.1101/2020.10.01.322834

**Authors:** Christiaan N. Hulleman, Rasmus Ø. Thorsen, Sjoerd Stallinga, Bernd Rieger

## Abstract

We have developed an engineered Point Spread Function (PSF) to enable the simultaneous estimation of dipole orientation, 3D position and degree of rotational constraint of single-molecule emitters from a single 2D focal plane. Besides giving access to orientation information, the Vortex PSF along with the vectorial PSF fitter avoids localization bias common in localization microscopy for fixed dipole emitters. We demonstrate this technique on reorienting single-molecules and using binding-activated localization microscopy on DNA intercalators, corroborating perpendicular azimuthal angles to the DNA axis for in-plane emitters but find a non-uniform distribution as a function of the polar angle. The Vortex PSF is realized by an affordable glass phase mask and has a compact footprint that can easily be combined with localization microscopy techniques on rotationally constrained emitters.

## Introduction

Single-Molecule Localization Microscopy (SMLM), with flavors like (F)PALM (1, 2), (d)STORM (3, 4) and (DNA)- PAINT (5, 6), have made nanoscale structural information beyond the diffraction limit more easily accessible to biologists. These super-resolution techniques commonly focus on localizing single emitters in the two dimensions of the focal plane, and sometimes in the third dimension along the optical axis. The role of molecular orientation in localization can often be ignored, as the fluorescent labels are flexibly attached to the biomolecule of interest and can rotate or wobble freely, thereby appearing as isotropic emitters. In case the fluorophores are more rigidly attached, emitter orientation is either a nuisance for estimating the position of the emitters accurately, or can give access to the orientation of the emitters. Imaging the orientation of constrained fluorescent labels has been used to visualize changes of fibroblasts under treatment (7) and reveal the underlying orientation of amyloid fibrils (8). Besides biological applications, the orientational information can also be used to visualize nanoscale deformations in material sciences (9). In this paper we demonstrate an engineered Point Spread Function (PSF), which allows in a unique manner the simultaneous estimation of *x, y* and *z* position of the emitter, as well as the polar and azimuthal angle *θ* and *ϕ* of the molecular orientation, and the degree of orientational constraint *g*_2_, all from a single 2D image. Thereby we can effectively combine orientation estimation and localization microscopy into a single process.

Visualization of emission patterns from fixed fluorescent emitters dates back to near-field studies (10) and studies with high-NA fluorescence microscopes, leading to the observation of ring-shaped spots originating from molecules oriented along the optical axis (11). Localization of these rotationally fixed molecules with a standard 2D Gaussian model leads to inaccuracies due to their dipole emission patterns (12). The localization accuracy of these fixed dipoles is significantly worse in the presence of aberrations (13), and with a small amount of defocus errors can easily amount to 100 nm (14). When defocusing fixed emitters, the observed pattern varies more as a function of orientation, opening up a way to estimate the orientation from a single image (15–20). A defocus of up to 1 *µm* spreads the emitted photons over many pixels, which has the drawback of adversely affecting localization precision. By interleaving in-focus localization with defocused spot fitting, a compromise between orientation estimation and localization precision can be made (21). The orientation can even be extracted from in focus single-molecule images in case of sufficiently high Signal-to-Noise Ratio (SNR) (22, 23).

The impact of rotational diffusion has been studied in ref. (24) in which the authors show that a localization bias on the order of 10 nm already occurs when the fluorophores are constrained to a cone half-angle of 60°. This localization bias can be avoided altogether by removing the radially polarized component (25, 26) or by neglecting the component responsible for bias (27), regrettably by doing so the SNR is reduced and the orientations that can be properly localized are limited.

Another inroad for measuring molecular orientation comes from fluorescence anisotropy measurements, which are done by alternating the excitation polarization and splitting the horizontally and vertically polarized emission to determine the rotational diffusivity (28, 29), or to distinguish different fluorophores (30). Though the anisotropy measurement can add rough orientation information to localization microscopy, it is impossible to differentiate ±45° azimuthal angles (7, 8, 31). By extending this method to more than two linear excitation polarizations the full in-plane orientation can be measured (32–34). Alas, this method does not give access to the out-of-plane angle. By scanning a radially polarized excitation focus the full orientation of the absorption dipole can be probed (35, 36), although this is not directly applicable to localization microscopy.

An alternative way to estimate orientation together with position comes from Point Spread Function (PSF) engineering, in which the phase profile of the emission beam is modified in the pupil plane, often in combination with polarization beam splitting approaches to separately measure different polarization components on the camera. Several techniques follow this principle, namely the double-helix PSF (37), quadrated pupil (38), bisected pupil (39) and tri-spot PSF (40), or based purely on polarization splitting and PSF shape fitting (41), where the PSF in each polarization channel changes as a function of the dipole orientation. Though these engineered PSFs encode the orientation well, the effective PSF size in each polarization channel is large, ranging from 4-16 times the Rayleigh criterion in size (*R* = 0.61*λ/*NA). This reduces the Signal-to-Background Ratio (SBR) (42, 43), and limits the density of emitters per frame for localization microscopy. In summary, fixed dipole emission compounded with additional aberrations and defocus leads to significant localization bias. Techniques that alleviate this bias often do not reveal the dipole orientation. The currently available techniques combining orientation and localization microscopy require a complex polarization beam splitting setup, or use a PSF design with a relatively large spot footprint on the camera.

In this paper we overcome these drawbacks by introducing the Vortex PSF, a PSF engineering approach to determine molecular orientation and position in all three spatial dimensions. The Vortex PSF does not require that different polarization components of the emission beam are imaged separately. The single imaged spot has a footprint of only 4-6 times the Rayleigh criterion in size, leading to a favorable SBR and high localization precision. Therefore the sparsity constraint for localization microscopy is on the same order of magnitude as a standard (non-engineered) PSF. The use of an aberration map, calibrated across the Field of View (FOV), with a fully vectorial PSF model in the parameter estimation avoids aberration induced biases and successfully reaches the Cramér–Rao Lower Bound (CRLB). This makes it possible to maintain precision and accuracy across standard FOVs of 10s of *µ*m. Furthermore the Vortex PSF enables access to the degree of rotational mobility, so that the flexibility of binding of the fluorophores can be probed as well. Implementation of the Vortex PSF into an existing setup is easy and can be realized for less than $4,000.

In the following we describe the optical principles of the Vortex PSF and the vectorial PSF fitting model with a calibrated aberration map. Simulations have been carried out to analyze the expected performance of the Vortex PSF for a range of parameter values. Proof-of-principle experiments have been conducted on isolated fixed single-molecules and the angles were verified by fitting defocused non-engineered PSFs. We showcase the method by tracking orientational jumps of single-molecules on a coverslip and imaging *λ*-DNA using Binding-activated localization microscopy (BALM) (44, 45).

## Methods

### Vortex PSF concept

The orientation of a constrained dipole emitter is characterized by three numbers, the in-plane azimuthal angle *ϕ*, the polar angle with respect to the optical axis *θ*, and a parameter quantifying the degree of rotational mobility *g*_2_ [Fig. 1(a)]. The *g*_2_ parameter is a ratio between the fixed and free dipole PSF which is sufficient to quantify the impact of orientational constraint, irrespective of the form of the constraint, e.g. “wobble-in-cone” or harmonic orientational potential well (46). A fixed dipole emitter that is oriented perpendicular to the optical axis (*θ* = 90°) emits fluorescence that is in-phase throughout the Fourier plane of the emission path. When imaged without the vortex, this emitter yields Gaussian-like spots on the camera due to constructive interference in the center. The emission from a dipole oriented along the optical axis (*θ* = 0°) captured by a high-NA objective has two regions on opposite sides of the Fourier plane of the emission path that are out of phase with respect to each other. Without the vortex this emitter yields a ring-shaped PSF with a zero in the middle due to destructive interference. Simulated images of standard PSFs show these PSF shapes close to in and out-of-plane orientations (*θ* = 80° and *θ* = 10°), yet the slight asymmetry arising from the azimuthal orientation cannot be identified by eye [Fig. 1(b)].

**Fig. 1.**
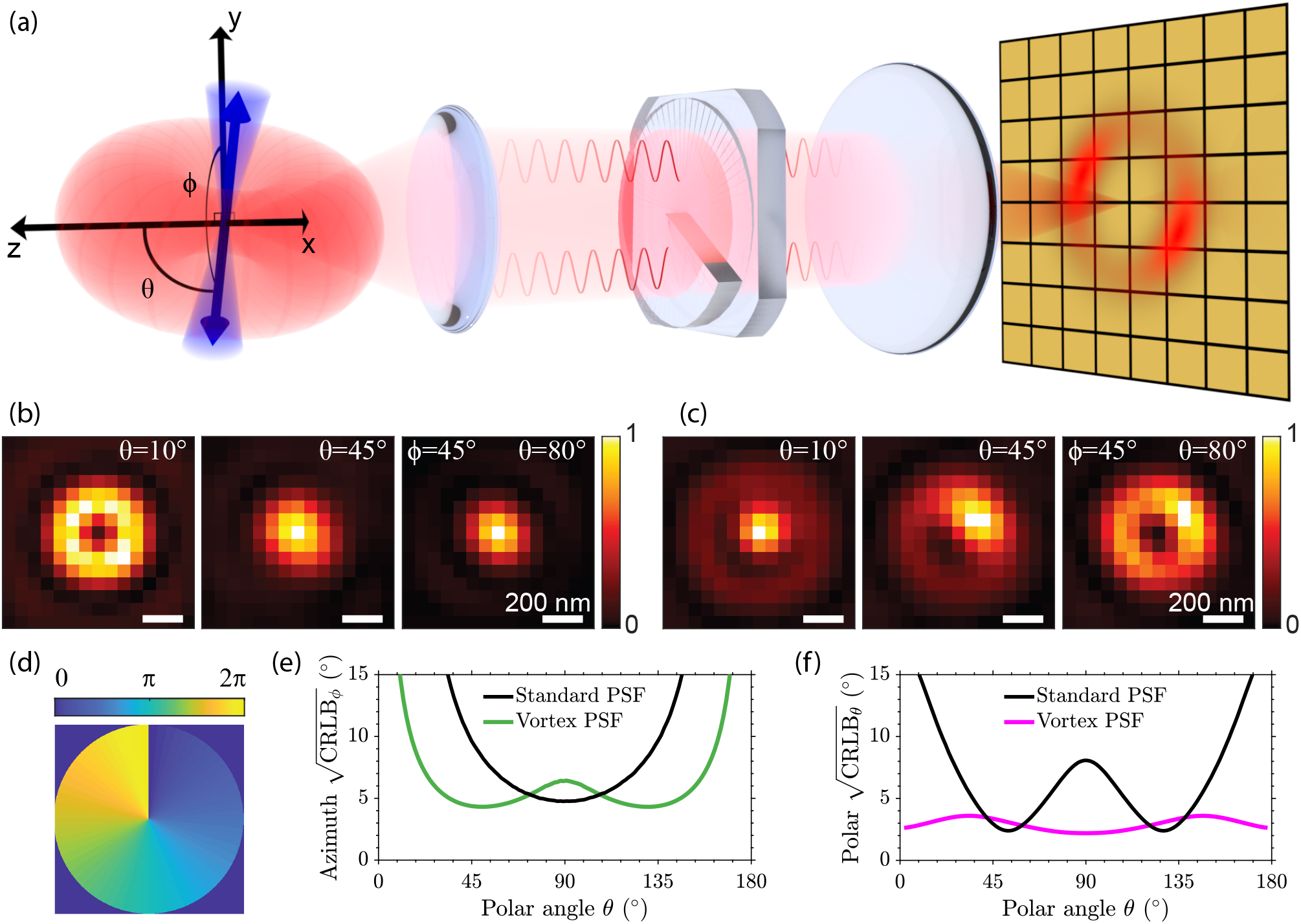
Vortex PSF concept. (a) A constrained dipole emitter is defined by the polar angle *θ* and azimuthal angle *ϕ*. The blue cone represents the degree of rotational constraint (*g*_2_) of the dipole emitter and the red torus-like shape represents the dipole emission. The microscope, equipped with a vortex phase plate, induces a radial *π* phase difference in the Fourier plane of the emission path. This results in an asymmetric donut-like shape for fixed emitters on the camera which we call the Vortex PSF. (b) Simulated standard PSF with polar angles from left to right (10, 45, 80 degrees) and an azimuthal angle of 45 degrees. (c) Simulated Vortex PSF with the same angles. (d) Phase profile of the vortex phase plate. (e) Azimuthal angle CRLB from simulated images as a function of the emitter polar angle (4000 signal photons and 10 background photons per pixel). (f) Polar angle CRLB as a function of the emitter polar angle with the same SBR.

The Vortex PSF can be realized as an addition to any standard fluorescence microscope and only requires a single phase plate in the Fourier plane of the emission path [Fig. 1(a)]. The phase plate consists of a single continuous phase vortex of topological charge *m* = 1. This is coincidentally also the lowest order component of a Double-helix PSF (47), and a discretized version of this spiral phase plate with only 3-4 phase-steps could be used to generate a rotating PSF (48). This phase plate shape is also used to create high-quality donut-shaped depletion and excitation profiles from Gaussian laser beams for STED (49) and MINFLUX (50). The phase delay of our phase plate is a spiral ramp from 0 to 2*π*, where radially opposing points always have a *π* phase difference between them [Fig. 1(d)].

Adding a vortex phase plate in the Fourier plane of the emission path inverts the phase relationship described earlier for a standard PSF and creates what we call the Vortex PSF. Now, out-of-plane (*θ* = 0°) orientations have a central spot and in-plane (*θ* = 90) orientations have a ring-shaped PSF. Due to the polarization and directional emission from the fixed dipole emitter, the intensity distribution changes along this ring as a function of the azimuthal angle. Simulated Vortex PSF shapes for polar angles (*θ* = 10°, *θ* = 45°, *θ* = 80°) indicate a substantial change as a function of the polar angle, as well as a clearly recognizable impact of the azimuthal angle (*ϕ* = 45°) on spot shape [Fig. 1(c)].

Fitting molecular dipole orientations using the standard infocus PSF is difficult because of its symmetries. The PSF is almost rotationally symmetric for all polar angles except around *θ* = 90°, where there is a slight asymmetry of the PSF as the spot is wider in one direction than the other. The azimuthal precision, quantified by the CRLB is indeed worse for all polar angles except near *θ* = 90° [Fig. 1(e)]. Furthermore, the standard PSF is also symmetric around the polar angles *θ* = 0° and *θ* = 90° yielding an unfavorable precision in estimating the polar angle around these angles [Fig. 1(f)]. This could explain why in-focus orientation estimations with a standard PSF have only been shown for polar angles 15° < *θ* < 70° (22, 23). These symmetries are broken by the Vortex PSF, resulting in a good precision over all possible orientations [Fig. 1(e,f)]. Of course, the precision of the azimuthal angle is still expected to diverge to infinity for polar angles approaching *θ* = 0° and *θ* = 180° as the azimuthal angle is undefined when the dipole is aligned along the optical axis. Note that due to the symmetry of the dipole it is sufficient to use half the unit sphere to uniquely define the dipole angle.

### Fitting model

We use standard Maximum Likelihood Estimation (MLE) using an image formation model that describes the expected photon count across the image as a function of the molecule position 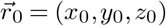, signal photon count *N* of the entire PSF on the camera, background photons per pixel *b*, and dipole orientation Ω_0_ = (*ϕ*_0_, *θ*_0_) with the degree of orientational constraint *g*_2_, giving a total of 8 parameters. The underlying PSF model is the fully vectorial PSF model, as in earlier work (13, 22, 51–53), but now extended to estimate the dipole orientation along with the degree of orientational constraint. The PSF model is taken to be the weighted sum of the freely rotating dipole PSF and the fixed dipole PSF corresponding to the equilibrium dipole orientation, as appropriate when the rotational diffusion is faster than the fluorescence lifetime (46):

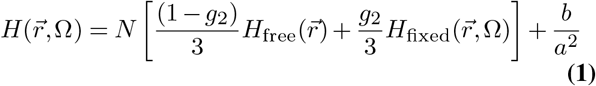

where *a* is the pixel size and 0 ≤ *g*_2_ ≤ 1 represents the degree of orientational constraint, with the limiting cases of a fully free dipole *g*_2_ = 0 and a fully fixed dipole *g*_2_ = 1. The vectorial model takes supercritical angle fluorescence (SAF) (54, 55) into account. Without accounting for SAF, emitters in the proximity of the coverslip could have an additional position bias up to ∼10 nm with a medium of water (*n* = 1.33) and ∼25 nm in air (*n* = 1). Further details on the image formation model are given in Supplement 1.

An important addition to the fitting model that we make is to take into account calibrated aberrations as done previously (52) and extended further to incorporate the field-dependence of aberrations (56). We improve upon this treatment by modeling the field dependence of the aberration coefficients using the so-called Nodal Aberration Theory (NAT) instead of 2D polynomials of arbitrary order. This approach is valid for optical imaging systems with small field angles (ratio of FOV to focal length), such as telescopes or microscopes, and has been devised by Shack and Thompson (57), and later extended and used in optical design and characterization studies (58–62).

The key prediction of NAT is that the dependence of aberration coefficients on the field coordinates is well approximated by Taylor series of a low order, such that there are specific relations between the coefficients of these series for different aberrations. Such relations exist for example between the Taylor series for the two astigmatic and the two coma aberration coefficients, giving rise to zero aberration loci (two, respectively, one for astigmatism and coma), so-called “nodes” in the FOV. The advantage of NAT in the current context is that the Taylor series fit for the different aberrations is more robust due to the predefined number of coefficients. Quantitative details on the aberration field dependence and the calibration procedure are given in Supplement 1. The aberration maps for 12 Zernike modes, determined from calibration measurements on 429 beads, are shown in Fig. S1 of Supplement 1. The NAT predictions are in excellent agreement for astigmatism and coma, and good for the other aberrations modes.

We have tested the model with extensive simulations to predict the experimental conditions under which the Vortex PSF can be used to correctly estimate the parameters Θ = (*x, y, z, N, b, ϕ, θ, g*_2_). We found that all model parameters can be estimated with precision at the CRLB for all molecular orientations and degrees of orientational constraint (Fig. S2 of Supplement 1), provided the signal-to-background ratio (SBR = *N/b*) is sufficiently high. A practical lower limit is around *SBR* ≥ 200 (Fig. S3 of Supplement 1). For dipole emitters orientated uniformly on unit sphere with *g*_2_ = 0.75 and SBR = 4000*/*10 the parameters can be estimated with a localization precision of *σ*_*xy*_ = 5.6 nm, *σ*_*z*_ = 27 nm and orientation precision of *σ*_*ϕ*_ = 5.5°, *σ*_*ϕ*_ = 3.1° and 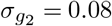. The polar precision appears almost constant over the unit sphere, whereas the azimuth precision performs well within polar angles (20° < *θ* < 160°) after diverging for emitters along the optical axis. Such polar range is notably broader than the standard PSF, as shown in Fig. 1e. The amount of rotational diffusion of an emitter affects the possible orientational and axial precision while it does not affect the lateral precision. The orientation can be estimated to a precision within *σ*_*θ*_ < *σ*_*ϕ*_ < 10° given a rotational diffusion *g*_2_ > 0.4. Outside this range the PSF becomes too smeared out, and the orientation information is mostly lost. The optimal axial performance is for fixed emitters *σ*_*z*_ = 23 nm (*g*_2_ = 1), whereas it worsens for freely rotating emitters up to *σ*_*z*_ = 49 nm (*g*_2_ = 0). In this case, when the emitters are freely rotating, the Vortex PSF has a slightly worse lateral precision compared to a non-engineered PSF (*σ*_*xy*_ = 5.9 nm versus *σ*_*xy*_ = 4.5 nm).

The estimation of all the parameters is uniform over a z-range of ±300 nm, with a region of interest of 15 × 15 pixels (Fig. S2 of Supplement 1). A larger region of interest should be used beyond ±300 nm to capture the whole PSF and attain reliable parameter estimates, though this is not optimal for dense single-molecule localization microscopy.

Simulations show furthermore that optical aberrations must be taken into account in the fitting model. Unknown or inaccurately calibrated aberrations with values deviating from the actual values by more than 36 m*λ* affect the imaging model so that the estimator introduces biases and no longer reaches the CRLB (Fig. S4 of Supplement 1). The aberration modes astigmatism and coma notably degrade the azimuthal precision by a factor of ∼ 2, and additionally, astigmatism also degrades the localization precision by a factor of ∼ 2. However, when the model is well calibrated with these aberrations, the estimator reaches the CRLB with no biases. This emphasizes the importance of using a good NAT aberration map to obtain reliable results over a large FOV.

### Experiments

The main component of the optical setup is a standard microscope (Ti-E, Nikon) with a 100x 1.49 NA objective (Fig. S5 of Supplement 1). A 4F relay system consisting of two 100 mm lenses (AC508-100-A-ML, Thorlabs) relays the original image plane of the microscope to the camera (Zyla 4.2 PLUS, Andor) and a vortex phase plate (V-593-10-1, vortex photonics) is placed in the Fourier plane between the two lenses. The vortex phase plate is mounted on a small XYZ stage (CXYZ05/M, Thorlabs) for alignment (see Supplement 1 and Fig. S6 of Supplement 1 for details on the alignment procedure), and a kinematic stage (KB75/M, Thorlabs) for quick removal and placement. The Vortex PSF should be used with a relatively narrow emission bandwidth up to ∼ 60 nm. The design wavelength of the phase plate does not need to exactly match the emission peak as long as both parameters are known and set correctly in the fitter.

When verifying the Vortex PSF’s functionality and performance, we imaged ATTO 565 embedded in a thin layer of PMMA (Polymethylmethacrylate) (Supplement 1). The PMMA polymer immobilizes the fluorophores and has a refractive index of *n* = 1.49 close to that of immersion oil (*n* = 1.518). Two z-stack acquisitions with a 100 nm step-size, 600 ms exposure and 300 W/cm^2^ epi-illumination are taken in quick succession with and without the vortex phase plate to compare the Vortex PSF to defocused orientation fitting.

To create a sparse sample of individual dipole emitters that are less constrained and can reorient over time, a low concentration of ATTO 565 without PMMA is spin-coated onto a coverslip. The resulting single-molecules are excited with 3 kW/cm^2^ in Total Internal Reflection Fluorescence (TIRF) conditions as the z-component of the TIRF field more effectively excites out-of-plane molecules. A relatively long exposure time of 900 ms is used to yield raw data with a very high SNR.

A wide variety of biological structures can be labeled with rotationally constrained fluorophores (21, 31, 33, 43). We chose to demonstrate our technique on frequently studied *λ*-DNA labeled with DNA intercalators (44, 45, 63) that transiently bind between the base pairs. The molecular dipole moment of the DNA intercalators is typically oriented perpendicular to the DNA axis (34, 64, 65), making this an ideal test case. For fixed dipole emission PSF fitting, the fluorophore must have a single molecular dipole moment, this makes bis-intercalators like TOTO (66) and YOYO (67) unsuitable. These dimeric fluorophores have two transition dipole moments between which the excitation energy can hop (68), resulting in emission from either of the two transition dipoles almost perpendicular to one another. We chose to use Sytox Orange which is believed to be a mono-intercalator (69) and further confirmed by elongation measurements matching that of mono-intercalators (63) (the exact chemical structure is undisclosed by the manufacturer).

In order to visualize the *λ*-DNA in a fluorescence microscope, molecular combing is used to align and stretch the DNA on a coverslip by a receding water-air interface (70, 71). For our sample preparation we adapted a spin-coating protocol (Supplement 1) that yields straight DNA strands with a variety of orientations (45). The sparsity required for localization microscopy is inherent from the transiently binding Sytox Orange that is essentially non-fluorescent when not intercalated (69). After applying a fresh batch of imaging buffer, 20,000 frames are acquired with a single frame exposure time of 100 ms. The sample is illuminated with 3 kW/cm^2^ circularly polarized total internal reflection excitation. This results in an excitation profile that is half in-plane and half out-of-plane, and a reduced background due to the limited penetration depth.

Details on the various sample preparation protocols for the different experiments are given in Supplement 1 along with details on the methods used for data analysis.

## Results

Figure 2 shows the results of a validation experiment for the Vortex PSF. Figure 2(a) shows through-focus images of a single-molecule in PMMA acquired in quick succession with and without the vortex phase plate. To verify the model over a large *z*-range a bigger region of interest (ROI) of 31 × 31 is used. These two imaging modes (with and without the phase plate) were used as input of two separate estimates. The first estimate serves as a ground truth measure and uses the standard PSF’s entire *z* stack to ensure high precision. This estimate differentiates from defocus imaging as described in the literature (16, 17, 21), where only a single focal slice is used. The second estimate uses the Vortex PSF, where an estimate is made for each focal slice. For the single-molecule in Fig. 2(a), its orientation found for the standard PSF *z*-stack fit (*ϕ, θ*) = (48°, 61°) agrees well with the angles found for the Vortex PSF *z*-slice fits (*ϕ, θ*) = (49 ± 1.5°, 61 ± 1.7°). Here the Vortex PSF uncertainty is the estimate of 11 focal slices, corresponding to a dynamic range of 1000 nm. Following the same procedure as for the molecule in Fig. 2(a), 21 different molecules’ estimated orientation is depicted in Fig. 2(b). The mean deviation between the orientation found with the standard PSF and Vortex PSF is (Δ*ϕ*, Δ*θ*) = (−0.4 ± 1.4°, −0.2 ± 1.2°), indicating no bias between the two imaging modes. The Vortex PSF’s mean precision is (*σ*_*ϕ*_, *σ*_*θ*_) = (2.3°, 1.8°), computed from the indicated error bars in Figure 2(b), which <u>c</u>omes close to the estimated lower limit 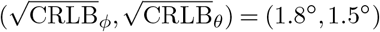. The estimated *z*-position shift between frames for the Vortex PSF of the molecule in Fig. 2(a) is 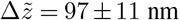, which matches the piezo shifts of 100 nm. The relationship between the estimated *z*-position and the piezo position averaged over 37 molecules is fitted with a linear function, resulting in a slope of −0.99 ± 0.01 and has a Root Mean Square Error (RMSE) of 16 nm over a 1200 nm range (see Fig. S7 of Supplement 1). To further show the quality of fit, cross-sections in the *x* − *z* and *y* − *z* planes are shown in Fig. 2(c)-(d). The agreement between the experimental data and fit with the vectorial Vortex PSF model is generally excellent. A striking detail is that even the fringe details away from focus match well. The lateral localization error, measured on individual *z*-slices between the two estimation modes, is 5 nm and 4 nm (RMSE) in the *x* and *y* direction. These validation experiments show that the orientation of fixed dipole emitters and their 3D position can be reliably estimated from individual focal-slice Vortex PSF images.

**Fig. 2.**
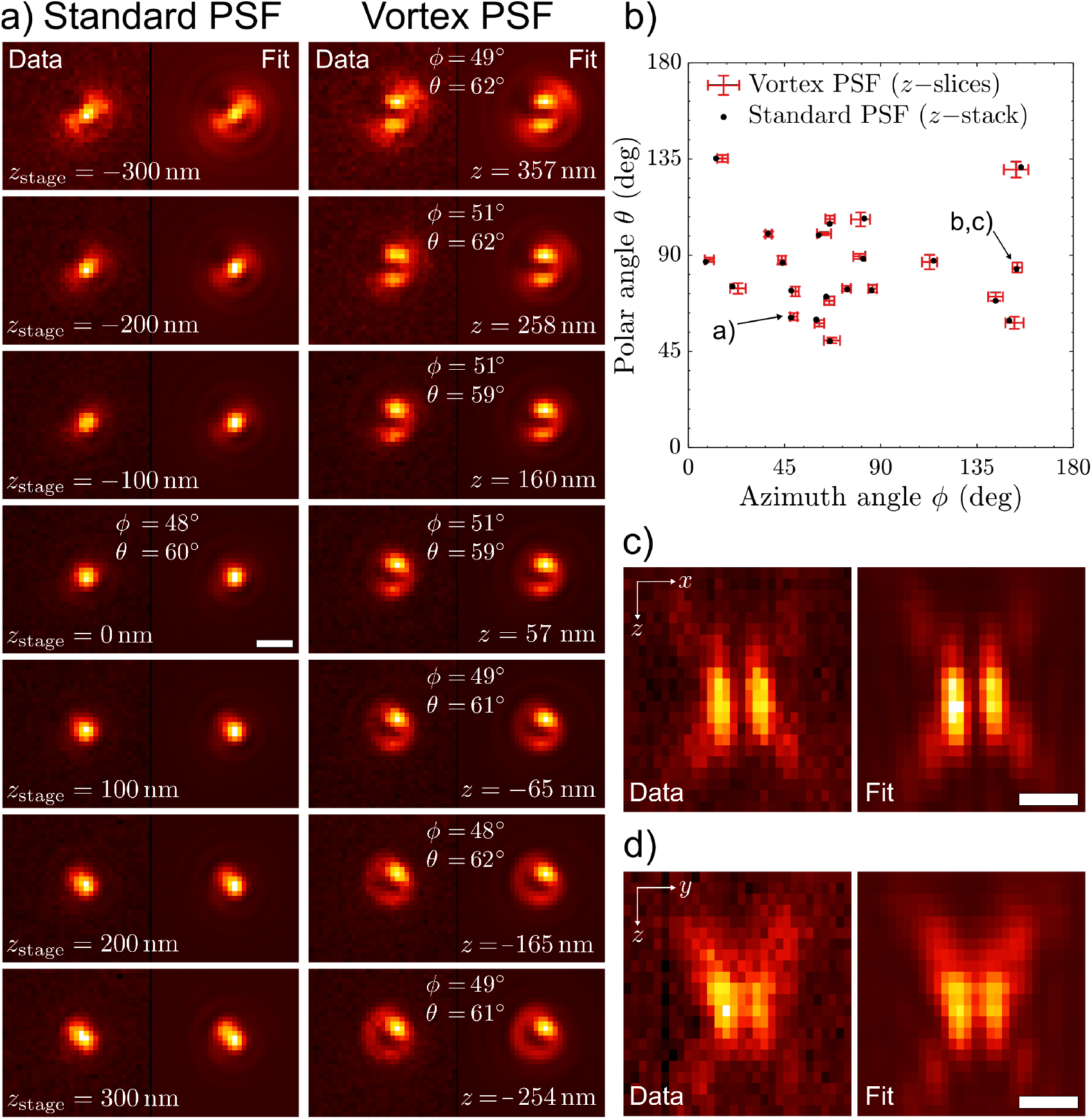
Vortex PSF validation experiment. (a) Standard PSF model fitted to an entire *z*-stack and the same molecule fitted for each *z*-slice independently with the Vortex PSF with its estimated parameters listed in each frame. All sub-image pairs are contrast stretched with the same factor for better visibility of the PSF shape. (b) Estimated orientations of 21 fixed molecules from a standard PSF z-stack and single-frame Vortex PSFs with error bars indicating one standard deviation. (c) The cross-section in *x* − *z* and (d) *y* − *z* of a Vortex PSF, with the measurement left and fit right (pixels are stretched proportionally in *z*). The estimated signal photon counts were in the range 4 − 50 × 10^3^, and the estimated background photon counts were in the range 10 − 40 photons per pixel. All scale bars are 500 nm.

The Vortex PSF can track dynamic changes in the orientation of single-molecules on a coverslip. We have observed that a portion of ATTO 565 single-molecules directly spincoated on glass show re-orientation when followed over time, indicative of metastable adhesion to the glass surface (see Visualization 1). The change in orientation of such a molecule can be directly seen from the dark region that shifts from the left to the right and back over 7 frames [Fig. 3(a)]. Time traces of 3 different molecules that show these re-orientation events indicate that the azimuthal angle changes, but the other parameters not that much [Fig. 3(b)]. These types of azimuthal angle changes would lead to large position biases in standard localization microscopy. The dip in the orientational constraint parameter *g*_2_ at a transition points to a more freely rotating molecule throughout that frame or a superposition of the orientation before and after the transition.

**Fig. 3.**
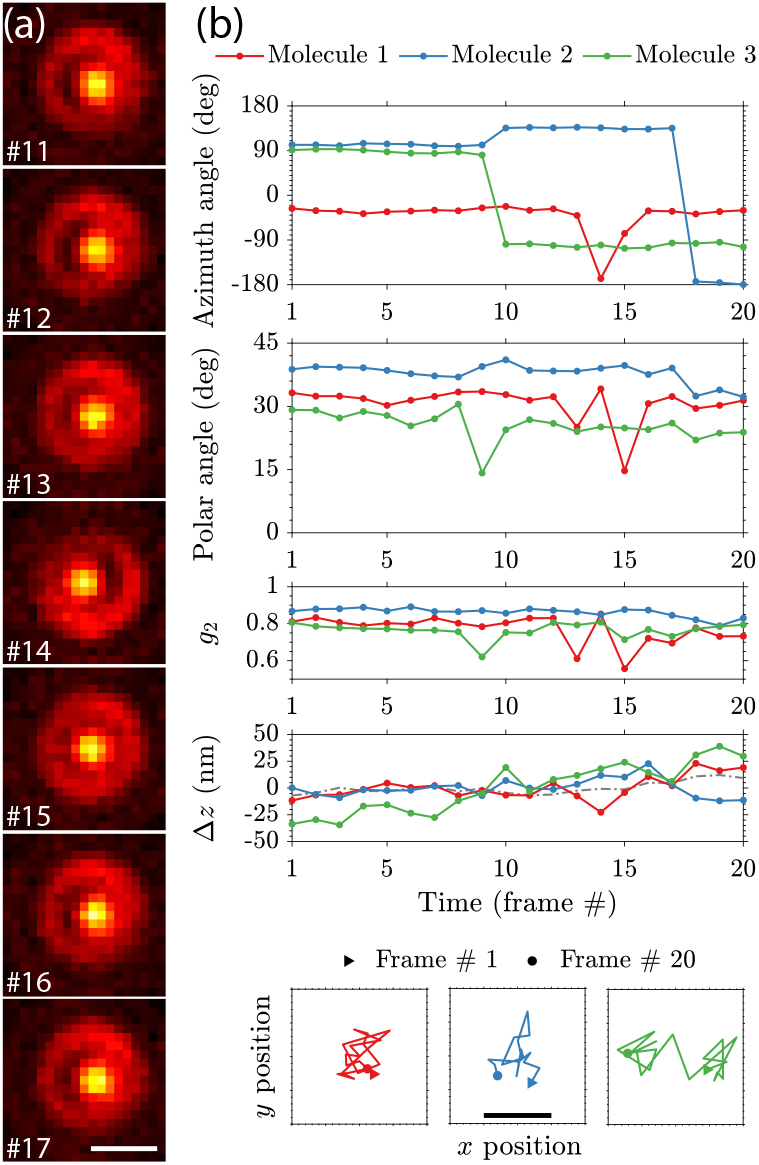
Re-orientation dynamics of single-molecules imaged with the Vortex PSF. (a) Consecutive raw Vortex PSF images of a molecule undergoing an orientational transition (900ms exposure, scale bar 500 nm). (b) Estimated parameters over 20 frames of three different molecules showing orientational transitions (molecule 1 shown in (a)). The gray curve in the Δ*z* plot shows the average upward drift (∼ 20 nm) from 10 stationary molecules. The *x*-*y* localization plots have 1 nm ticks and a scale bar of 10 nm.

Figure 4 shows the potential of combining localization microscopy and orientation estimation by imaging *λ*-DNA. The super-resolution reconstruction can be color-coded with one of the orientation parameters (azimuthal or polar angle, degree of orientational constraint), or the *z*-position. Fig. 4(a) shows a color-coding with the azimuthal angle. The in-plane molecular orientation is clearly perpendicular to the orientation of the DNA-strands. Analyzing a single strand shows an azimuthal angle difference between the fluorophore and the DNA axis of 84° with a median absolute deviation (MAD) of 20° [Fig. 4(b)]. This is essentially the same as found before (Δ*ϕ* = 87° and MAD(Δ*ϕ*) = 18°) (34) and similar to measurements with YOYO (64, 65). The degree of orientational constraint *g*_2_ is estimated with a peak at *g*_2_ = 0.8, which corresponds to a maximum angle *α* ∼ 31° in the framework of the wobble-in-cone model [Fig. 4(c)], which is slightly larger than ∼ 22° found previously (34, 64). All these parameters are estimated while attaining a lateral resolution (35 nm FWHM *λ*-DNA line-width) typical for BALM (44, 45) [Fig. 4(d)]. If the field-dependent aberrations were not taken into account in the fitting model, the localization distribution becomes non-Gaussian with a *λ*-DNA line-width twice as broad (73 nm FWHM in Fig. S8 of Supplement 1).

**Fig. 4.**
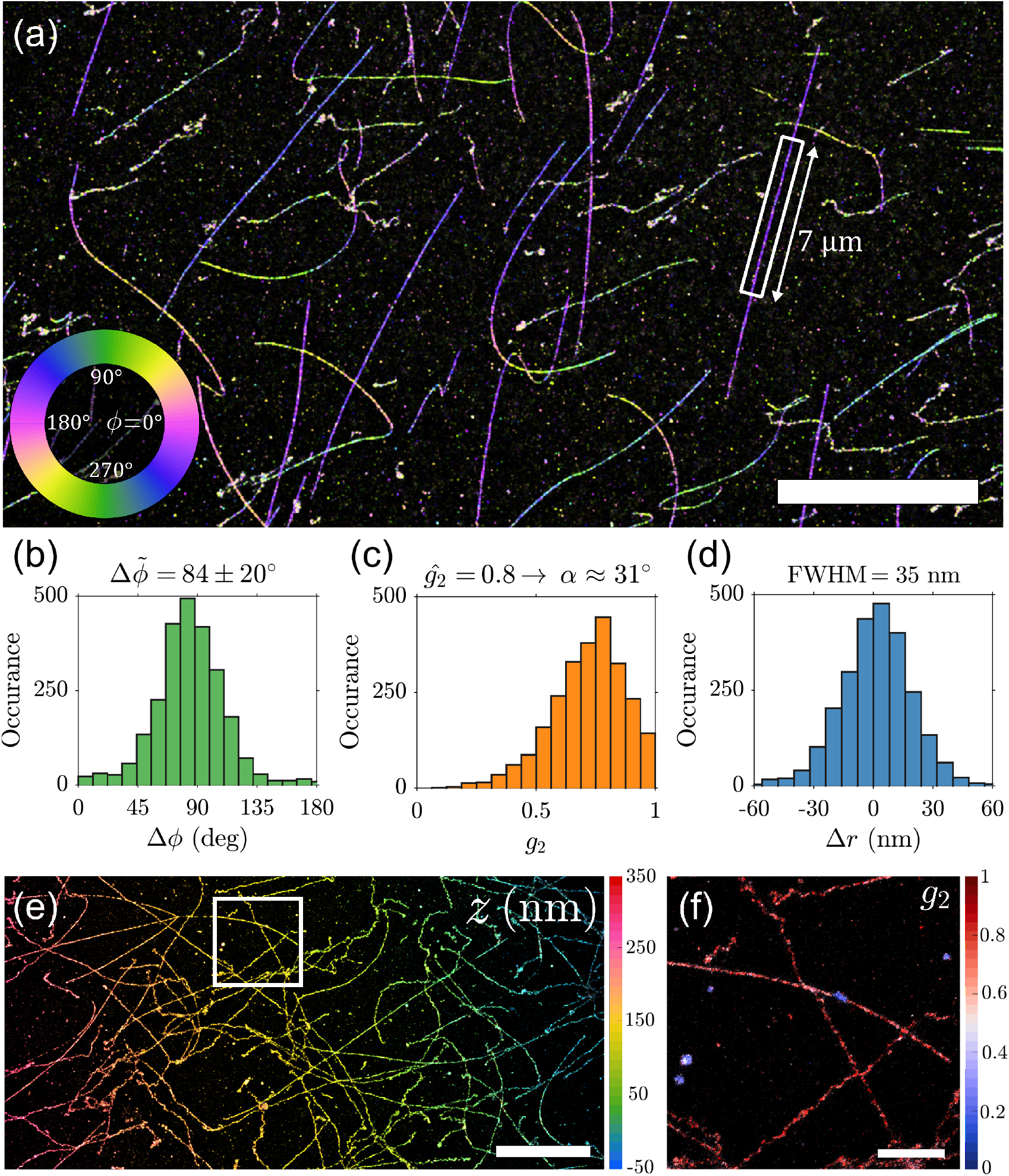
Super-resolution image of *λ*-DNA. (a) *λ*-DNA colorized as a function of the azimuthal angle. (b) Azimuthal angle histogram relative to the DNA axis from the DNA strand highlighted in (a) (Δ*ϕ* = 84°, MAD(Δ*ϕ*) = 20°). (c) Distribution of *g*_2_ from the same strand, the fitted peak *g*_2_ = 0.8 corresponds to a wobble cone angle *α* = 31°. (d) Position deviation from the spline fit to the DNA axis with a FWHM of 35 nm (Filtered to 45° ≤ *θ* ≤ 135°). (e) Tilted *λ*-DNA sample colorized as a function of z-position. (f) *λ*-DNA from highlighted region in (e), colorized as a function of *g*_2_. Scale bars are (a) 10 *µ*m, (e) 5 *µ*m, (f) 1 *µ*m.

Furthermore, we use an intentionally tilted *λ*-DNA sample to demonstrate the ability to resolve the lateral structure independent of defocus. This sample is shown in Fig. 4(e), where the estimated *z* position reveals the slope of the tilted *λ*-DNA sample. Fig. 4(f) shows that color-coding with the degree of orientational constraint *g*_2_ can be used to identify patches where binding to partially detached DNA strands and/or non-specific binding occurs. These are visible with a near free dipole value *g*_2_ ∼ 0.3, as opposed to the nearly fixed dipole value *g*_2_ ∼ 0.8 of the DNA strands. The achieved precision from these experiments, determined from repeated localizations of multiple on events from the same emitter (Fig. S9 of Supplement 1), is 5.4 nm and 29.7 nm in the lateral and axial dimension, whereas the azimuth and polar angle precision are 6.0 deg and 3.9 deg, respectively. The localization and orientation precision values determined in this way come close to the predicted CRLB and are within the estimated error bars.

The orientational binding landscape can be investigated by analyzing correlations between the orientational parameters (*θ, ϕ, g*_2_), in combination with probing with different excitation polarization. To that end we have imaged the same region both with and without a quarter waveplate (QWP) in the illumination laser path [Fig. 5(a)]. Without the QWP the excitation polarization is vertical and in-plane, most effectively exciting molecules around *θ* = 90° [Fig. 5(b)]. With QWP the polarization is half in-plane and half out-of-plane resulting in less selective excitation. This results in a polar distribution with more localizations around *θ* = ±40° [Fig. 5(c)]. Without the QWP we find the expected relative angle of Δ*ϕ* = 87° between the fluorophore and DNA axes with almost no dependence on the polar angle [Fig. 5(b)]. With the addition of the QWP a correlation between the polar and azimuthal angle becomes visible. Next to the population of molecules with close to in-plane orientations, that still have the expected relative angle Δ*ϕ* = 84° (green histogram), a second population of molecules with more out of plane angles *θ* = ±40° appears, that has a relative angle of Δ*ϕ* = 65° (gray histogram). Figs. 5(d) and (e) show that there is also a correlation between the polar angle and the degree of orientational constraint, where the population of molecules with near in-plane orientation have a smaller *g*_2_ value compared to the population of molecules with clear out-of-plane orientation, showing a looser orientation for the in-plane population. These correlations between the orientational parameters are only found in the *λ*-DNA experiments and are independent of the DNA orientation. Control experiments on fixed single-molecules in air and PMMA show a uniform distribution over the azimuthal angles with no correlation to polar angle or degree of orientational constraint (see Fig. S10 of Supplement 1).

**Fig. 5.**
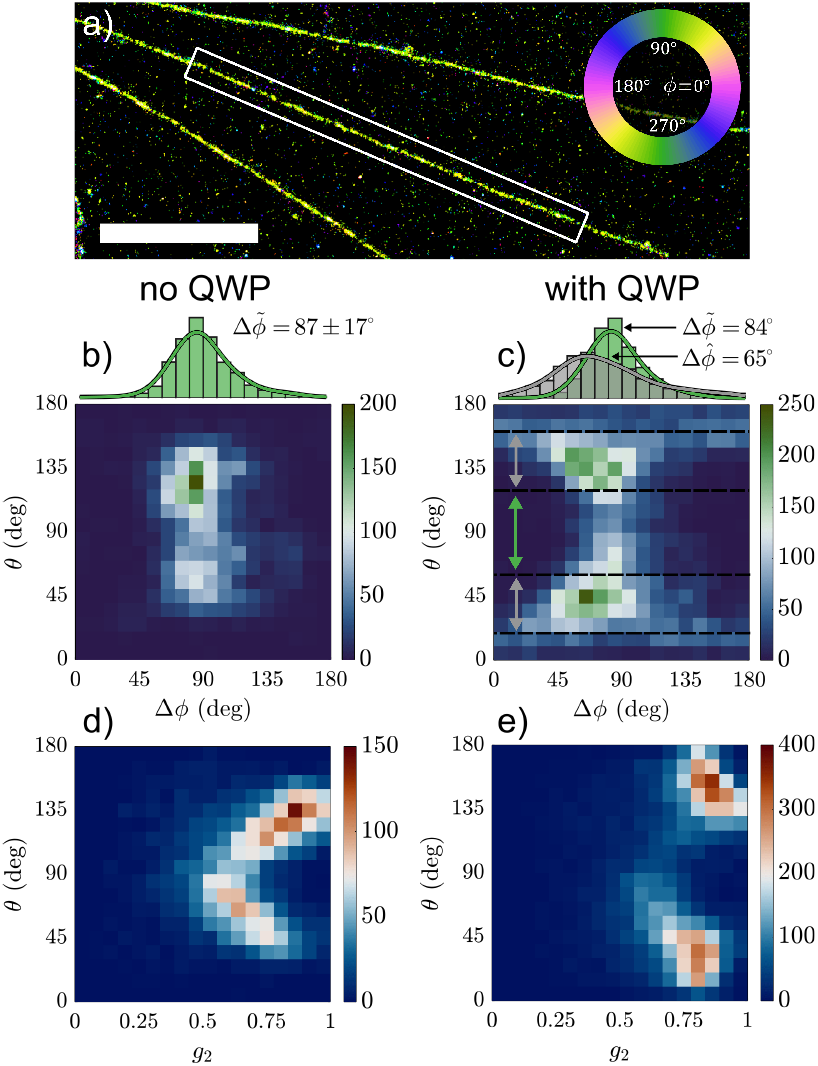
Correlation between orientation parameters of *λ*-DNA under different excitation polarization. (a) Super-resolution image of *λ*-DNA strands color coded with the same azimuthal map as Figure 4(a). Scale bar is 3 *µ*m. (b) Histogram of the relative azimuthal angle with respect to the DNA axis from data-set without QWP in the illumination path. (c) 2D histogram of polar angle vs relative azimuthal angle with the QWP and two highlighted regions of polar angles around *θ* = 90° (green) and *θ* = ±40° (gray). (d) 2D histogram of polar angle vs rotational diffusion coefficient *g*_2_ without QWP. (e) 2D histogram of polar angle vs rotational diffusion coefficient *g*_2_ with QWP.

One would expect a uniform polar angle distribution for free DNA strands as Sytox Orange should intercalate in any orientation, averaging out any base pair selectivity along the strand. The proximity of the coverslip to the DNA strand could limit the physical space available for intercalators, thereby creating a non-uniform polar angle distribution and possibly shift the equilibrium azimuthal angle away from the orientation perpendicular to the DNA strand. However, this does not explain the preference for Δ*ϕ* = 65° over Δ*ϕ* = 115° for both polar angle regions. This may originate from the helical structure of the DNA and the binding potentials between the intercalator and the DNA.

A different hypothesis for the observed correlations between the orientational parameters is a change of the DNA structure to S-DNA due to overstretching. Previously it was found that overstretching results in a change of azimuthal angle to Δ*ϕ* ∼ 54° (64), comparable to the value we find. The typical length of *λ*-DNA strands in our datasets is 17.4 *µ*m which is 7.4% longer than its crystallographic length (72). Although this corresponds to a relatively low percentage of overstretching it is possible that the binding affinity is not as low in the proximity of the coverslip as for free DNA strands, resulting in a relatively large population of tilted orientations compared to the DNA strand. We did not observe a correlation between the orientational parameters and the position along the DNA strand, that would correspond to domains along the strand with different orientational binding. This makes this hypothesis less likely.

## Conclusion

With the Vortex PSF we achieve a fit that is both accurate, avoiding the large position biases commonly seen with fixed dipole emitters in localization microscopy, and precise, close to the CRLB. A key ingredient is that we properly take into account calibrated field-dependent aberrations and supercritical angle fluorescence in the vectorial PSF model to avoid these biases. The relatively compact spot shape enables a more favorable trade-off between the precision of estimating the position and orientational parameters compared to other PSF designs.

We have visualized and measured orientational transitions of single-molecules with metastable attachment to a glass surface. Furthermore we have applied our method to *λ*-DNA, corroborating previous findings that the azimuthal angle of the intercalator dipoles is almost perpendicular to the DNA axis. Our method uncovered a preferential polar orientation of intercalators attached to *λ*-DNA on a coverslip, along with a correlation between orientation and orientational constraint. We have applied the Vortex PSF to these two cases to illustrate its functionality, but in principle, the Vortex PSF can be applied to any sparse sample of constrained dipole emitters. Combined with a sparsity inducing single-molecule localization microscopy technique, super-resolution images can be complemented with orientation information to differentiate various sub-sets in the data, such as identifying different binding modes, different orientational configurations or local deformations. Nanoscale interactions could be investigated using chemical models on the single-molecule scale.

An interesting question to address in future studies would be to compare the Vortex PSF to fundamental (quantum) limits of the estimation of orientational parameters (73, 74). Another step to advance the Vortex PSF concept could be a speed up of the fitting algorithm by developing a GPU implementation, or by using spline interpolated models obtained from PSF calibrations (75) extended to take into account field-dependent aberrations. The feasibility of estimating the emission wavelength in addition to the other parameters could also be investigated. Finally, the analysis could be extended into the regime of slow orientational diffusion. In that regime the illumination polarization has an impact on PSF shape, implying that modulation of the illumination polarization into the method could generate useful information on the orientational constraint and diffusion of the molecule.

## Funding

European Research Council (ERC) (648580); National Institutes of Health (NIH) (U01EB021238).

## ACKNOWLEDGEMENTS

We thank Srividya Ganapathy for help preparing the pH 5.5 solution and Marijn Siemons for help with the vectorial PSF fitter applied to SAF conditions.

## Disclosures

The authors declare no conflicts of interest.

See Supplement 1 for supporting content (supplemental figures, materials and methods, and calculations). The code is available here (github.com/imphys/vecfitcpu_vortex) and the data here (10.4121/c.5136125).

## Notes

https://github.com/imphys/vecfitcpu_vortex

https://doi.org/10.4121/c.5136125

### Competing Interest Statement

The authors have declared no competing interest.

