## Supplement 1 for "Simultaneous orientation and 3D localization microscopy with a Vortex point spread function"

**This document provides supplementary information to "Simultaneous orientation and 3D localization microscopy with a Vortex point spread function".**

**localization microscopy | fixed dipole emission | PSF-engineering | super-resolution | orientation estimation | fluorescence | Lambda DNA**

### 1. Imaging PSF model

We use a full vectorial PSF model in the vortex PSF fitting, as initially described in (1), and extend it to incorporate varying degrees of orientational constraint and orientational diffusion (2). The expected photon count at pixel $l$ depends on the molecule position $\vec{r}_0 = (x_0, y_0, z_0)$, the signal photon count $N$, background photons per pixel $b$, and the molecular orientation $\Omega_0 = (\phi_0, \theta_0)$ together with the degree of orientational constraint $g_2$, giving a total of 8 parameters. The integration of the PSF model $H$ gives the expected photon count:

$$\mu_l = \int_{D_l} \mathrm{d}x\mathrm{d}y\, H(\vec{r} - \vec{r}_0, \Omega) \tag{S1}$$

where the integration is over the pixel area $D_l$ of size $a \times a$. The PSF model is taken to be the weighted sum of the free dipole PSF and the orientation-dependent fixed dipole PSF, where the relative weights are determined by $g_2$. This can be written as

$$H(\vec{r}, \Omega) = N\left[\frac{(1-g_2)}{3} H_{\mathrm{free}}(\vec{r}) + \frac{g_2}{3} H_{\mathrm{fixed}}(\vec{r}, \Omega)\right] + \frac{b}{a^2} \tag{S2}$$

The PSF of a fixed dipole emitter is

$$H_{\mathrm{fixed}}(\vec{r}, \Omega) = \sum_{i,j=x,y,z} A_{ij}(\vec{r}) d_i(\Omega) d_j(\Omega) \tag{S3}$$

where $d_i(\Omega)$ are the components of the dipole unit vector $\vec{d}(\Omega) = (\sin\theta\cos\phi, \sin\theta\sin\phi, \cos\theta)$. The average PSF of a freely rotating dipole emitter is

$$H_{\mathrm{free}}(\vec{r}) = \frac{1}{3} \sum_{i=x,y,z} A_{ii}(\vec{r}) \tag{S4}$$

with

$$A_{ij}(\vec{r}) = \sum_{k=x,y} w_{ki}(\vec{r}) w^*_{kj}(\vec{r}) \tag{S5}$$

where the functions $w(\vec{r})$ represent the electric field component in the image plane proportional to the emission dipole component $j$. These functions can be expressed as integrals over the pupil plane:

$$w_{kj}(\vec{r}) = \frac{1}{\pi} \int \mathrm{d}^2\rho\, A(\vec{\rho}) \exp\left(\frac{2\pi i W(\vec{\rho})}{\lambda}\right) q_{kj}(\vec{\rho}) \exp(-i\vec{k}\cdot\vec{r}) \tag{S6}$$

where the integration is over normalized pupil coordinates $\vec{\rho}$, $A(\vec{\rho})$ is the aplanatic amplitude correction factor, and $q_{kj}(\vec{\rho})$ are the polarization vectors given in full detail in (3). The wavevector $\vec{k}(\vec{\rho})$ depends on the normalized pupil coordinates by

$$\vec{k}(\rho) = \frac{2\pi}{\lambda}\left(\mathrm{NA}\rho_x, \mathrm{NA}\rho_y, \sqrt{n^2 - \mathrm{NA}^2\vec{\rho}}\right) \tag{S7}$$

with $n$ the refractive index of the medium. The aberration function $W(\vec{\rho})$ describes the zone function of the vortex phase plate: $K(\vec{\rho}) = \beta/(2\pi)$ where $\beta = \arctan(\rho_x/\rho_y)$ is the azimuth pupil coordinate, and further includes field-dependent aberrations as described in section 2.

The imaging model's partial derivatives with respect to the parameters are needed for the MLE optimization routine. These are easy to evaluate for the signal photon count $N$, background photons per pixel $b$, and diffusion weights $g_2$ as these appear linear in the imaging model $\mu_k$. The derivatives with respect to the fit parameters $\Theta = (x, y, z, \phi, \theta)$ are similar to (4) but now slightly more elaborate:

$$\frac{\partial \mu_l}{\partial \Theta} = N\left[\frac{1-g_2}{3} \sum_{k=x,y} \sum_{j=x,y,z} \int \mathrm{d}^2\rho\, 2\Re\left\{w^*_{kj} \frac{\partial w_{kj}}{\partial \Theta}\right\} + \frac{g_2}{3} \sum_{k=x,y} \sum_{j=x,y,z} \int \mathrm{d}^2\rho\, 2\Re\left\{w^*_{kj} d_j \frac{\partial (w_{kj} d_j)}{\partial \Theta}\right\}\right] \tag{S8}$$

where $w_{kj} = w_{kj}(r - r_0)$ and $d_j = d_j(\Omega)$. The derivatives of $w_{jk}$ with respect to the coordinates of the emitter are needed here:

$$\frac{\partial w_{kj}(\vec{r} - \vec{r}_0)}{\partial \vec{r}_0} = \frac{i}{\pi} \int \mathrm{d}^2\rho\, A(\vec{\rho}) \exp\left(\frac{2\pi i W(\vec{\rho})}{\lambda}\right) q_{kj}(\vec{\rho}) \times \vec{k}(\vec{\rho}) \exp(-i\vec{k}(\vec{\rho})\cdot(\vec{r} - \vec{r}_0)) \tag{S9}$$

as are the derivatives of the dipole vector $\vec{d}(\Omega)$ with respect to the polar and azimuthal angles:

$$\frac{\partial \vec{d}(\Omega)}{\partial \phi} = (-\sin\theta\sin\phi, \sin\theta\cos\phi, 0) \tag{S10}$$

$$\frac{\partial \vec{d}(\Omega)}{\partial \theta} = (\cos\theta\cos\phi, \cos\theta\sin\phi, -\sin\theta). \tag{S11}$$

### 2. Field dependent aberration coefficients

The aberrations function $W(\vec{\rho})$ is conventionally expressed as a linear sum of the root mean square (RMS) normalized Zernike polynomials $Z_n^m(\vec{\rho})$: $W(\vec{\rho}) = \sum_{n,m} A_n^m Z_n^m(\vec{\rho})$. In most studies the Zernike coefficients are constant parameters. In this study we follow the approach of (5) and (6) by taking the dependence on the position in the FOV into account, i.e. we take the Zernike coefficients to be functions of the field coordinates $(x,y)$. These functions $A_n^m(x,y)$ are determined from a calibration procedure. We make a through-focus image stack of a set of beads randomly spread over the FOV, as opposed to the procedure of (6), where a single bead is positioned on a series of grid positions by the microscope stage. For each bead the Zernike coefficients are retrieved using our previous method (7). According to Nodal Aberration Theory (NAT), the aberration coefficients $A_n^m(x,y)$ can be suitably described by low order Taylor series in $x$ and $y$:

$$A_n^m(x,y) = \sum_{jk} \gamma_{nmjk} x^j y^k. \tag{S12}$$

The set of coefficients $\gamma_{nmjk}$ of these Taylor series for different positions are related (8), which we use to our advantage as this decreases the number of parameters to be determined from experiment. The NAT-model is fitted to the measured $A_n^m$ at the beads' positions by a straightforward least-squares fit. With this calibration procedure, the estimated Zernike coefficients can effectively be interpolated over the entire imaging field.

We take into account Zernike modes with $n + |m| \leq 6$, which include primary and secondary astigmatism, coma, and spherical aberration, and trefoil, and use the analysis of (9), where polynomials in the field coordinates up to order $6-n$ are used in the NAT description of the field dependence of the contributing Zernike modes.

In the following, these expressions are summarized as implemented in our fitting using a set of perturbation coefficients $(\chi, \xi, \delta, \mu, \eta, \kappa, \nu)$. Primary astigmatism with perturbation coefficients $\chi$ is given by:

$$\begin{aligned} A_2^{-2} = {} & \chi_1(x^3y + xy^3) + \chi_2(x^2y + y^3) + \chi_3(x^3 + xy^2) \\ & + \chi_4(x^2+y^2) + \chi_6(xy^2 - x^3) + \chi_7 2xy^2 \\ & + \chi_8 y - \chi_9 x + \chi_{10}(2xy^2) + \chi_{11} x + \chi_{12} y + \chi_{13} \end{aligned} \tag{S13}$$

$$\begin{aligned} A_2^{2} = {} & \chi_1(y^4 - x^4) - \chi_2(x^3 + xy^2) + \chi_3(x^2y + y^3) \\ & + \chi_5(x^2+y^2) - \chi_6 2x^2y + \chi_7(y^3 - x^2y) \\ & + \chi_8 x + \chi_9 y + \chi_{10}(y^2 - x^2) + \chi_{11} y - \chi_{12} x + \chi_{14}, \end{aligned} \tag{S14}$$

primary coma with coefficients $\xi$ by:

$$\begin{aligned} A_3^{-1} = {} & \xi_1(x^3 + xy^2) + \xi_2 x^2 + \xi_3 xy + \xi_4 x + \xi_5(x^2+y^2) \\ & + \xi_7 y + \xi_8 x + \xi_9 \end{aligned} \tag{S15}$$

$$\begin{aligned} A_3^{1} = {} & \xi_1(y^3 + x^2y) + \xi_2 xy + \xi_3 y^2 + \xi_4 y + \xi_6(x^2+y^2) \\ & + \xi_7 x - \xi_8 y + \xi_{10} \end{aligned} \tag{S16}$$

and primary spherical aberration with coefficients $\delta$ by:

$$A_4^0 = \delta_1(x^2+y^2) + \delta_2 x + \delta_3 y + \delta_4. \tag{S17}$$

The next aberration order trefoil with coefficients $\mu$ is given by:

$$\begin{aligned} A_3^{-3} = {} & \mu_1(3y^2x - x^3) + \mu_2(y^2 - x^2) \\ & + \mu_3 2xy + \mu_4 x + \mu_5 y + \mu_6 \end{aligned} \tag{S18}$$

$$\begin{aligned} A_3^{3} = {} & \mu_1(y^3 - 3x^2y) - \mu_2 2xy + \mu_3(y^2 - x^2) \\ & + \mu_4 y - \mu_5 x + \mu_7 \end{aligned} \tag{S19}$$

secondary astigmatism with coefficients $\eta$ by:

$$A_4^{-2} = \eta_1 2xy + \eta_2 y + \eta_3 x + \eta_4 \tag{S20}$$

$$A_4^{2} = \eta_1(y^2 - x^2) - \eta_2 x + \eta_3 y + \eta_5 \tag{S21}$$

secondary coma with coefficients $\kappa$ by:

$$A_5^{-1} = \kappa_1 y + \kappa_2 \tag{S22}$$

$$A_5^{1} = \kappa_1 x + \kappa_3 \tag{S23}$$

and finally we include secondary spherical aberration, which is expected to be constant in the included polynomial order but here modeled to match (S17), with coefficient $\nu$:

$$A_6^0 = \nu_1(x^2+y^2) + \nu_2 x + \nu_3 y + \nu_4. \tag{S24}$$

The independent perturbation coefficients are determined using least-squares over all bead measurements to relate the perturbation coefficient to the Zernike coefficients for the field coordinates. Once the perturbation coefficients are known, the equations (S13 – S24) are used as predictors based on the detected molecule position. The retrieved field-dependent Zernike aberration coefficients for our microscope are shown in Fig. S1.

### 3. Simulation setup

Simulated point spread functions (PSFs) are generated according to the vectorial PSF model described in section 1. The NA is taken to be 1.45, the wavelength 597.5 nm, the refractive index of the imaging medium 1.33, coverslip 1.523, immersion medium 1.518, and a pixel size of 65 nm in object space in a region of interest (ROI) of 15x15 pixels. Unless stated otherwise, we take 4000 detected signal photons on the camera and 10 background photons per pixel, and we neglect readout noise but add Poisson noise to each image. The number of photons corresponds to the number of photons captured into the NA and thus spread over the entire FOV. The fraction of signal photons captured within the ROI is 0.44

and 0.46 for the standard and vortex PSF, respectively. The simulations are run for 10,000 randomized instances with coordinates taken from a uniform distribution over $\pm 1$ pixel and molecular dipole orientations uniformly distributed on the unit sphere. That is, if $u$ is a uniform random number from the distribution $U[0,1]$, the simulated angles are taken to be $\phi_0 = \pi u$ and $\theta_0 = \arccos(1-2u)$.

### 4. Vortex Alignment

A 1:1 optical relay is built to place the vortex phase plate in the Fourier plane of the emission path of the microscope as illustrated in Fig. S5. The phase plate is placed roughly halfway between the two relay lenses. To align the vortex phase plate, defocused images are taken of 1 $\mu$m beads (TetraSpeck Fluorescent Microspheres Size Kit, ThermoFisher). If the vortex phase plate is aligned properly, all the PSFs should have the same shape throughout the FOV as shown in Fig. S6(b). When the vortex phase plate is not in the correct axial position the PSF will vary over the field of view. This is because light from different areas in the sample do not pass through the vortex phase plate in the same place as it is not in the Fourier plane. The main parts of the PSF to observe for alignment is the peak in the center that does not move when the phase plate is moved and a ring that moves with the phase plate. The process of aligning the vortex phase plate involves moving the rings to be centered over the peaks over the entire FOV. The vortex phase plate should be translated along the optical axis until all the beads look the same throughout the field of view (same offset between the peak and the ring throughout the FOV) [Fig. S6(c)-(d)]. Thereafter the vortex phase plate can be shifted horizontally and vertically so the dark spot overlaps with the center of the bead [Fig. S6(e)-(f)]. Lastly the beads are translated along the optical axis and defocused in the opposite direction to verify that the rings also overlap with the center of the bead there. If the rotation direction of the vortex phase plate is unknown ($K(\vec{\rho}) = \beta/(2\pi)$ or $K(\vec{\rho}) = -\beta/(2\pi)$), then fixed single molecules can be fitted with both orientations and the correct setting will have a visually better fit (should be especially evident on molecules with $\theta = \pm 45°$).

### 5. Sample Preparation

Cover slips (22x22 mm No. 1.5, Marienfeld-Superior) and microscope slides (Microscope slides, Menzel Gläser Thermo Scientific) are cleaned by sonication in ethanol for 15 minutes and are then blown dry with nitrogen. All further mentions of cleaned cover slips and microscope slides are cleaned the same way except the cover slips for the $\lambda$-DNA samples. The microscope slides for the $\lambda$-DNA samples have a 5-10 mm hole drilled in them in advance to make it possible for the imaging medium to be added from above. Lambda DNA ($\lambda$-DNA) (Lambda DNA, Thermo Scientific) is aliquoted into 10 $\mu$L portions in PCR tubes and stored at -20 °C. Ascorbic acid (Ascorbic acid, Merck) is divided into ~3 mg portions in PCR tubes and the mass written on the tubes and stored at 4 °C. To make a pH 5.5 solution, 1 $\mu$L of 400 mM HCl and 600 mM Tris, is diluted in Milli-Q (MQ) water to a pH of 5.5 (approximately 90 mL). Part of a 5 mM stock solution of Sytox orange (SYTOX Orange Nucleic Acid Stain, Invitrogen) is diluted in TE buffer (Tris-EDTA buffer solution pH 7.4, Supelco) by 4 tenfold steps to 500 nM and stored at 4 °C.

Fixed single molecule samples are made by sparsely embedding ATTO 565 in a thin layer of PMMA. 100 mg of PMMA (Poly(methyl methacrylate), Sigma-Aldrich) is dissolved in 10 grams of Toluene (Toluene, Sigma-Aldrich). ATTO 565 (ATTO 565, Sigma-Aldrich) is diluted in MQ water in 100 fold steps to $\sim 5$ $\mu$M. The ATTO 565 dilution is further diluted in PMMA/Toluene in 100 fold steps to a $\sim 5$ pM concentration. A 20 $\mu$L droplet of the mixture is placed on a clean cover slip in the spin-coater and is spun at 3000 RPM for 2 minutes. Two strips of double sided tape (Permanent Double Sided Tape, Scotch) are placed ~1.5 cm apart on a cleaned microscope slide and the cover slip is placed on top with the PMMA facing the tape side.

For single molecules without PMMA the same procedure is followed except now ATTO 565 is diluted only in MQ water to a final concentration of $\sim 500$ pM. After spin-coating the coverslip is placed, coated side down, on double sided tape on a microscope slide.

The $\lambda$-DNA samples are prepared by dropping a $\lambda$-DNA solution onto a rotating silanized cover slip (10, 11). These cover slips are cleaned more extensively by sonication for 1 hour each in ethanol, acetone and then ethanol again. The cleaned cover slips are stored in ethanol. Before silanizing the surface they are removed from the ethanol and blown dry with nitrogen. An individual dry cover slip is then placed in 15 mL of Poly-L-lysine solution (Poly-L-lysine solution 0.01% sterile-filtered, Sigma-Aldrich) for 5 minutes and slightly shaken 2-3 times. Thereafter the silanized cover slip is rinsed with MQ water and left to dry overnight. A 10 $\mu$L $\lambda-$DNA aliquot is thawed and 990 $\mu$L of the pH 5.5 solution is added. For a silanized cover slip the optimal combing pH appears to be just below pH 5.5 (12). 40 $\mu$L of this solution is applied in a drop wise fashion to the silanized cover slip on the spin-coater rotating at 2500 RPM for 30 seconds. Thereafter the speed is increased to 7000 RPM and 5 mL of MQ water is applied to rinse away non attached DNA and left spinning for 2 minutes to dry. A square hole is cut into a piece of double sided tape (64621, Tesa) and placed around the pre-drilled hole in the microscope slide. The cover slip is placed on the tape and pressed down with the $\lambda$-DNA side towards the tape. The ascorbic acid is hydrated with TE buffer to a concentration of 200 mM just before the experiment. 25 $\mu$L of the ascorbic acid dilution is mixed with 5 $\mu$L of 500 nM Sytox Orange and 470 $\mu$L of TE buffer. 200 $\mu$L of the imaging buffer with a final concentration of 5 nM Sytox Orange and 10mM ascorbic acid in TE buffer is added to the sample through the hole in the microscope slide while on the microscope, focusing and imaging starts as soon as possible.

To determine the aberration maps, 180 nm orange bead samples were used. A 1/100 dilution of 180 nm orange beads (PS-Speck Microscope Point Source Kit, ThermoFisher) is made by mixing 10 $\mu$L of beads with 990 $\mu$L of MQ water. Using 10 $\mu$L of this dilution, 7-10 small droplets are placed around the centre of the cleaned cover slip and allowed to dry for ~3 hours or overnight. Two strips of double sided tape (Permanent Double Sided Tape, Scotch) are placed ~1.5 cm apart on a cleaned microscope slide and the cover slip is placed on top with the beads facing the tape side. When ready to image a 20-30 $\mu$L droplet of the mounting medium (in our case immersion oil) is placed on the edge between the cover slip and microscope slide. The capillary action gradually distributes the mounting medium between the cover slip and the microscope slide. The sample is placed on the microscope when the mounting medium has reached the other side.

### 6. Data analysis

The acquired images are offset and gain corrected to convert analog-to-digital units (ADUs) into photon numbers (13). Then, candidate pixels with a single molecule signal are identified using an intensity threshold typically chosen as the background plus a constant of around 10. These candidate pixels are segmented into ROIs of size 15x15 pixels centered at the local centroid. The local centroid gives a better first estimate of the emitter's position than the local maximum due to the Vortex PSF shape. These ROIs are fitted with a vectorial PSF model using Maximum Likelihood estimation (MLE). The vectorial PSF model incorporates emitter parameters described in section 1 and is further tailored to take field-dependent aberrations into account described in section 2. The optical parameters and experimental settings are set to match the experimental setup in Fig. S5.

The resulting localizations are corrected for sample drift following the method of Schnitzbauer et al. (14), implemented in the Picasso software (v0.2.8), available at github.com/jungmannlab/picasso. The lateral drift for the $\lambda$-DNA experiments was on the order of 1.5 pixels ($\sim$ 100 nm) over 30 minutes. Subsequent localizations of the same emitter during the emitter's on-time are linked under the condition that the position and orientation estimate between subsequent localizations is less than 3 times the largest uncertainty as shown in Fig. S9.

All images are rendered with a Gaussian blurring using the scripts from the INSPR toolbox (15). The images in Fig. 4 are rendered with a super-resolution pixel size of 6.5 nm, and image in Fig. 5 with a pixel size of 20 nm. Estimating the axes of single DNA strands is performed by fitting a spline to the localization data. First, a DNA strand is selected from the localization data incorporating all localizations. A spline curve is fitted to the localizations using MATLAB's built-in function *fit()* employing a smoothing spline with a smoothing parameter of $10^{-1}$. In order to determine the azimuthal orientations of molecules with respect to the DNA axis, the shortest distance between a given molecule and a point on the spline curve is determined, and the tangent line to the point on the spline is calculated using finite differences, giving the local DNA-strand orientation.

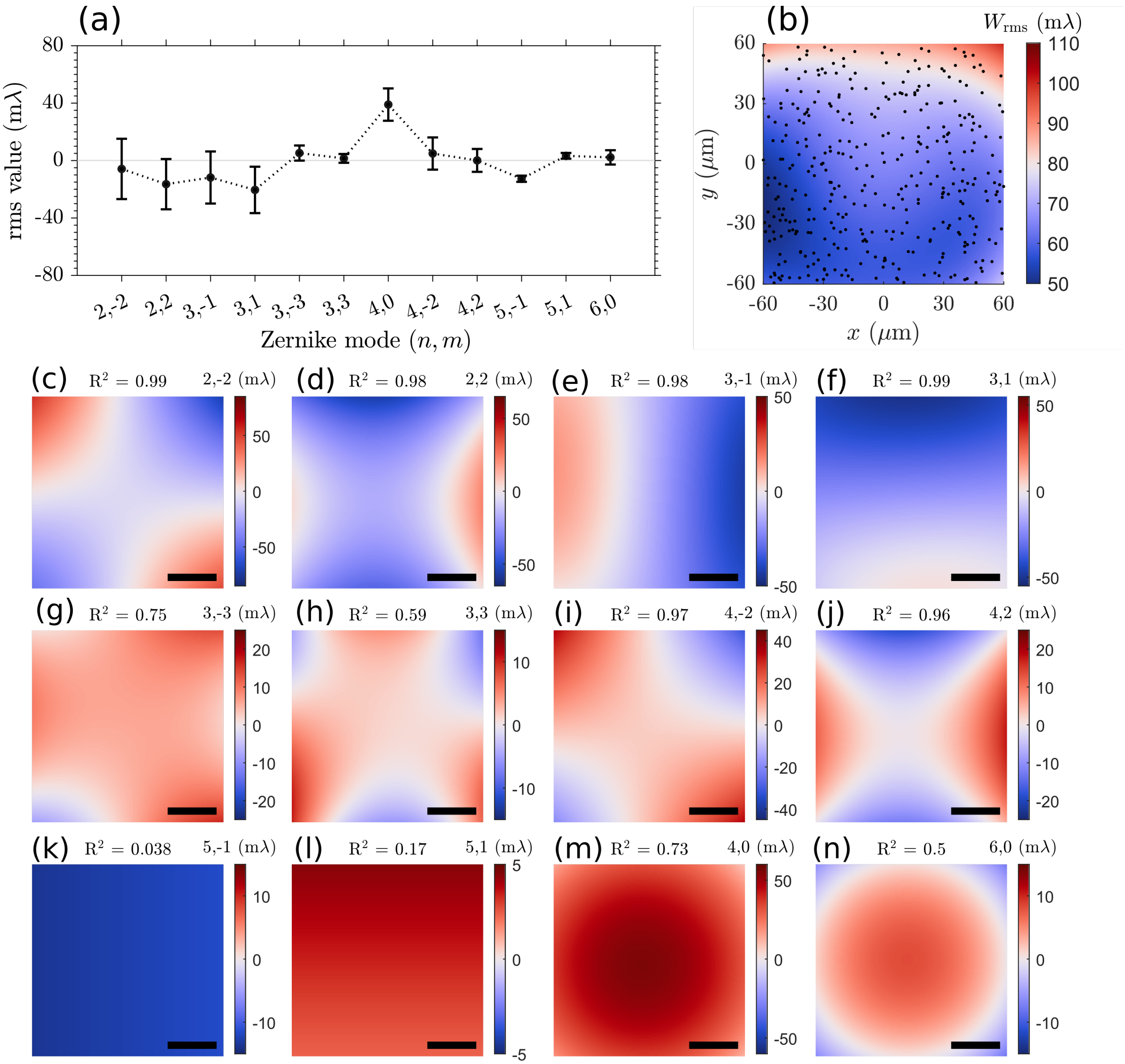


**Fig. S1.** Quantification of aberration retrieval and correction in the field of view (FOV). (a) Fitted Zernike modes and retrieved aberration coefficients over the entire FOV. The coefficients are averaged over 429 bead localizations with error bars indicating the mean and one standard deviation. (b) The total wavefront error from the field aberration surfaces in (c-n). The black dots indicate individual bead locations. (c-n) Fitted field aberrations from the coefficients in (a) and Zernike surfaces as presented in section 2, with $R^2$ as the quality of fit. Scale bars in (c-n) are 30 $\mu$m.

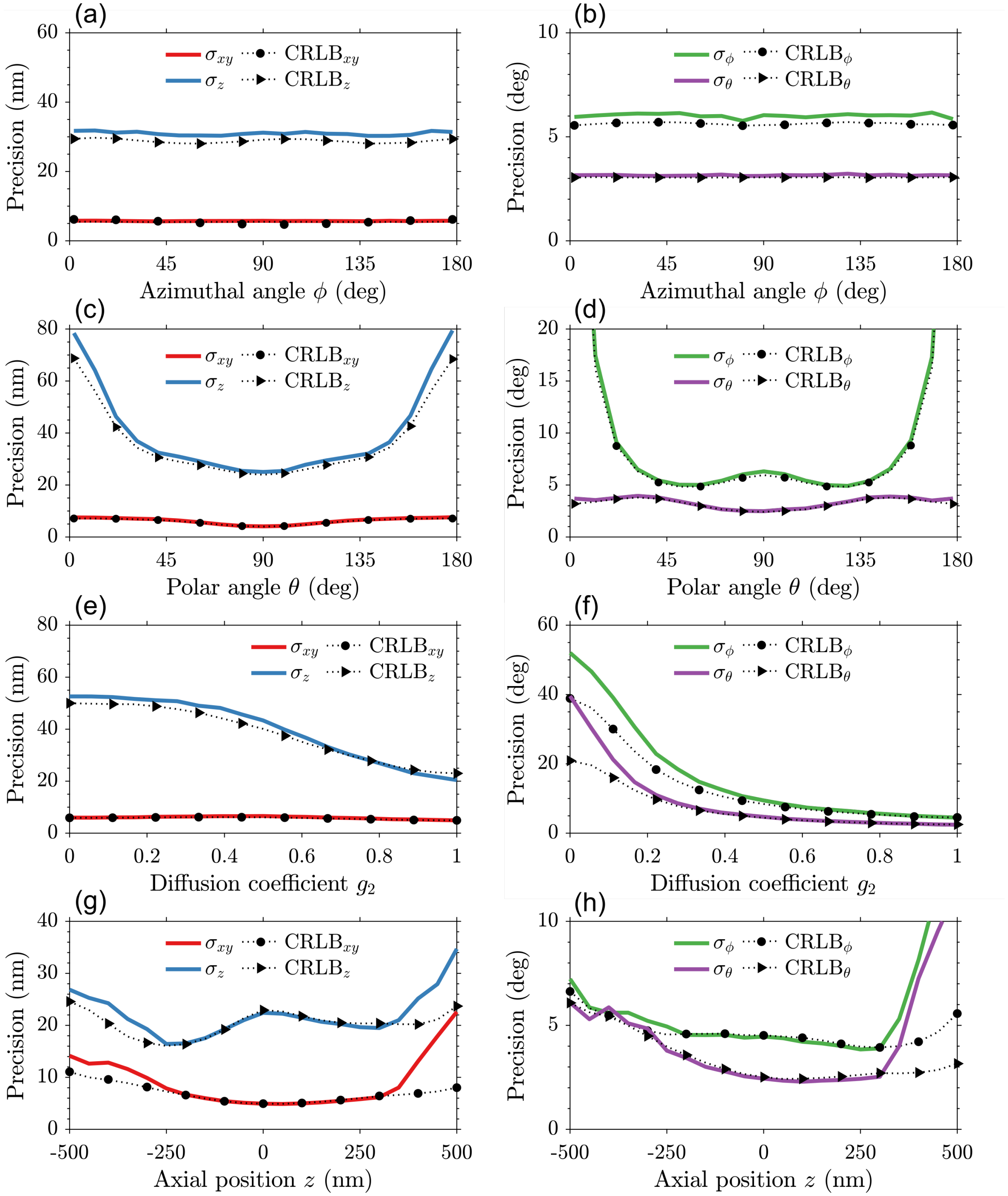


**Fig. S2.** Simulation study of the impact of molecule orientation, rotation diffusion, and axial position on the precision. (a) Average lateral and axial localization precision and (b) orientation precision as a function of the molecule's azimuthal angle with its polar angle uniformly chosen on a sphere, and $g_2 = 0.75$. (c-d) Localization and orientation precision as a function of the polar angle with its azimuthal angle uniformly chosen on a sphere ($g_2 = 0.75$). (e-f) Precision as a function of diffusion coefficient with the molecule's orientation uniformly chosen on a sphere. (g-h) Localization and orientation precision as a function of emitter's axial position, again with the molecule's orientation uniformly chosen on a sphere ($g_2 = 0.75$). The estimator's performance (solid colorized lines) is at the CRLB (black dashed lines with symbols) for all molecule orientations, regardless of the rotational diffusion. The estimator achieves the CRLB for axial positions of $|z| < 300$ and starts to diverge outside this region as the Vortex PSF footprint becomes too large to be contained within the $\mathrm{ROI} = 15 \times 15$. Simulation parameters as described in section 3.

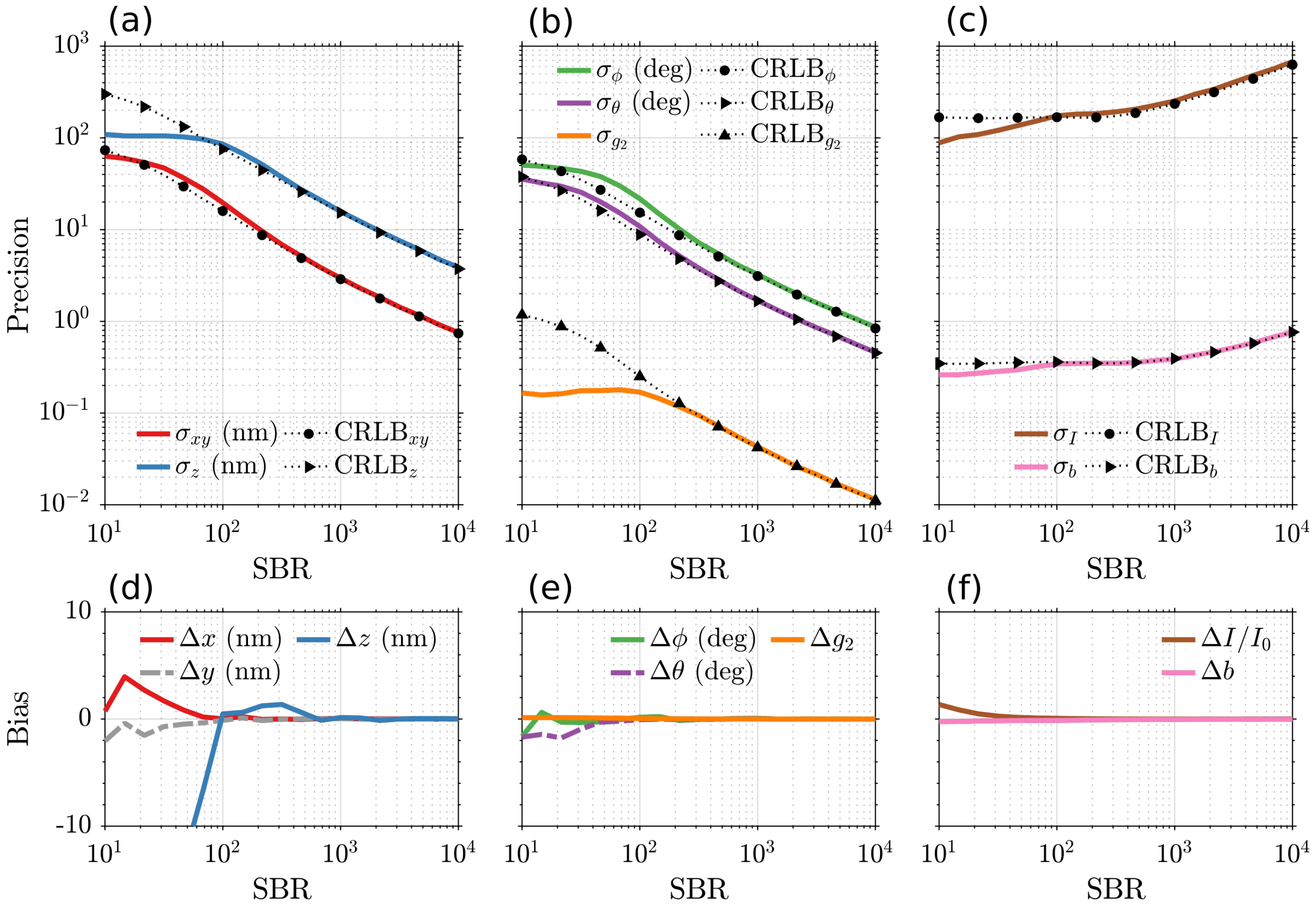


**Fig. S3.** Simulation study of the impact of signal to background ratio ($\mathrm{SBR} = I/b$) on precision and bias of the estimated parameters. (a) Average lateral and axial localization precision using a vortex PSF model on simulated emitters, simulated with the vectorial PSF (see Methods) as a function of SBR. (b) The orientation precision of the azimuthal and polar angles together with the diffusion coefficient. (c) The photon precision of the signal photons and background photons per pixel. The estimation performance is at the CRLB except at SBR levels below $10^2$, indicating that the number of signal photons is too low to assess the vortex PSF in the image. (d-f) Similarly, the bias of the estimated parameters compared to ground truth for the (d) localization, (e) orientation (f), and photon errors. Simulation parameters as described in section 3.

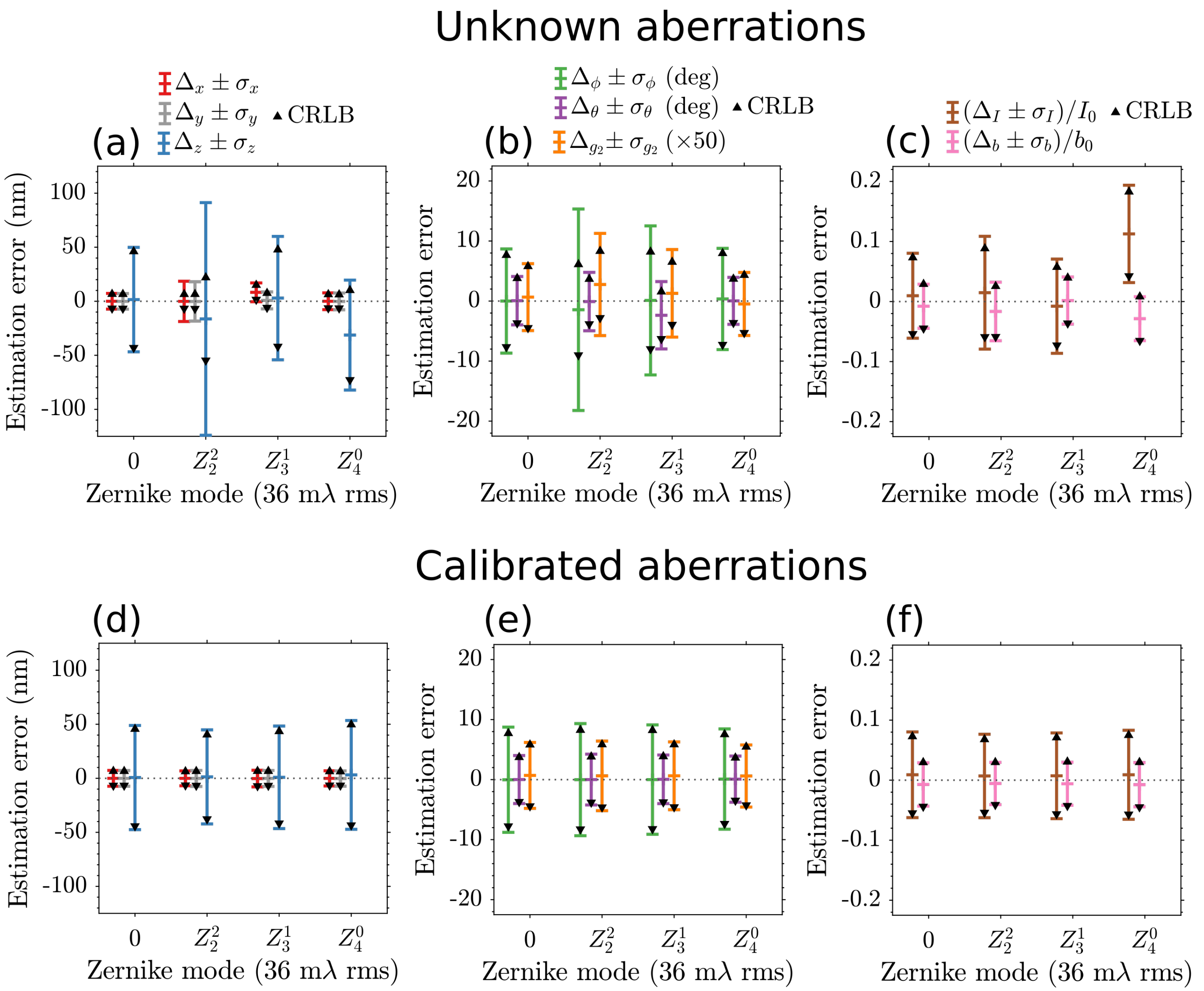


**Fig. S4.** Simulation study of the impact of unknown and known single-mode aberrations fitting with the vortex PSF. (a) Lateral and axial localization error using an unaberrated vortex PSF model on simulated emitters with single-mode aberrations: first-order astigmatism $Z_2^2$, first-order coma $Z_3^1$, and first-order spherical $Z_4^0$ with RMS value of 36 m$\lambda$. The error bars indicate the mean and one standard deviation with the black marker indicating the CRLB. (b) In the same way, the orientation error and (c) signal and background photon error. (d-f) Same as (a-c) but including the aberrations in the Vortex PSF model. Fitting with various 36 m$\lambda$ known aberrations the performance is at the CRLB, and the bias is removed.

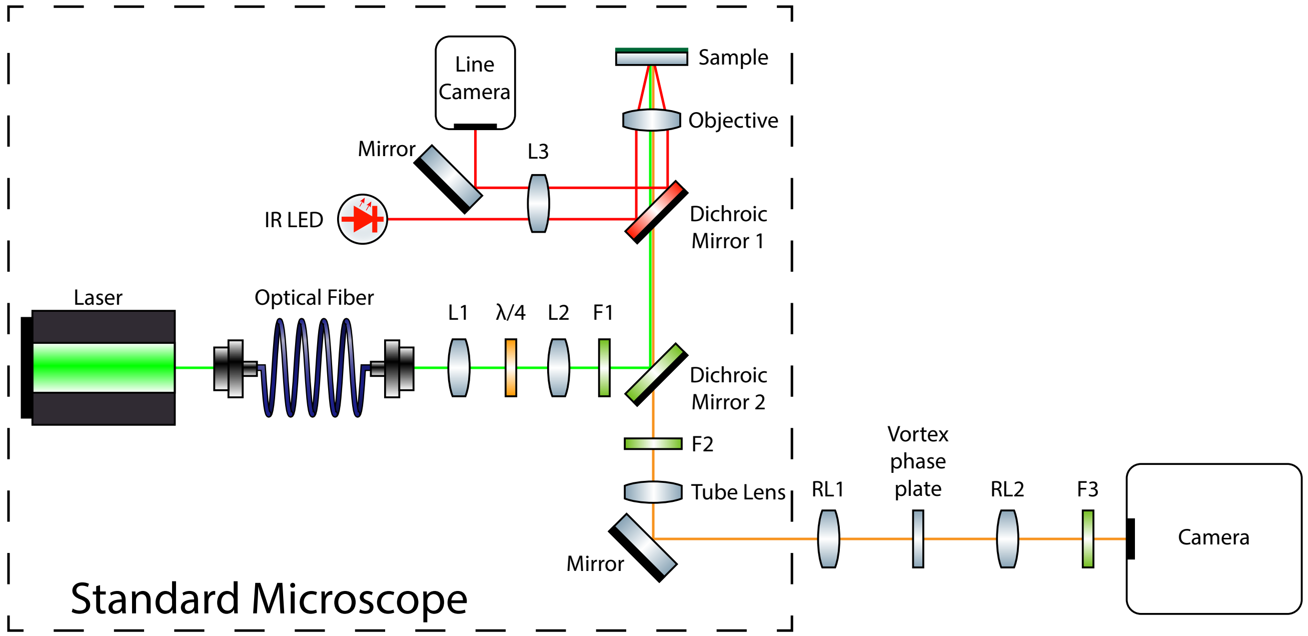


**Fig. S5.** The optical setup consists of a simple addition to a standard microscope (Ti-E, Nikon). Lenses RL1 (AC508-100-A-ML, Thorlabs) and RL2 (AC508-100-A-ML, Thorlabs) relay the original image plane to the camera (Zyla 4.2 PLUS, Andor) with no additional magnification. The Fourier plane is placed in between these two lenses, where the Vortex phase plate (V-593-10-1, vortex photonics) is placed. The standard TIRF microscope has a focus lock consisting of an infrared light emitting-diode, offset lens L3, Dichroic mirror 1 and a line camera (any unspecified components are part of the Nikon Ti-E or its accessories). The excitation laser (Sapphire 561-150 CW, Coherent) is coupled into a fiber and collimated by lens L1 and thereafter focused onto the back focal plane of the objective (CFI Apochromat TIRF 100XC Oil, Nikon) with lens L2. By translating the fiber face the excitation beam angle coming out of the objective can be adjusted to total internal reflection conditions in the sample. The $\lambda/4$ waveplate converts the linearly polarized laser beam to circular polarization, and the excitation spectrum is filtered by F1 (ZET405/488/561/640x, Chroma). Dichroic mirror 2 (ZT405/488/561/640rpc, Chroma) splits the excitation and emission path and the emission spectrum is further filtered by F2 (ZET405/488/561/640m-TRF, Chroma) and F3 (FF01-609/57-25, Semrock). The tube lens focuses the image from the sample to the front focal plane of lens RL1.

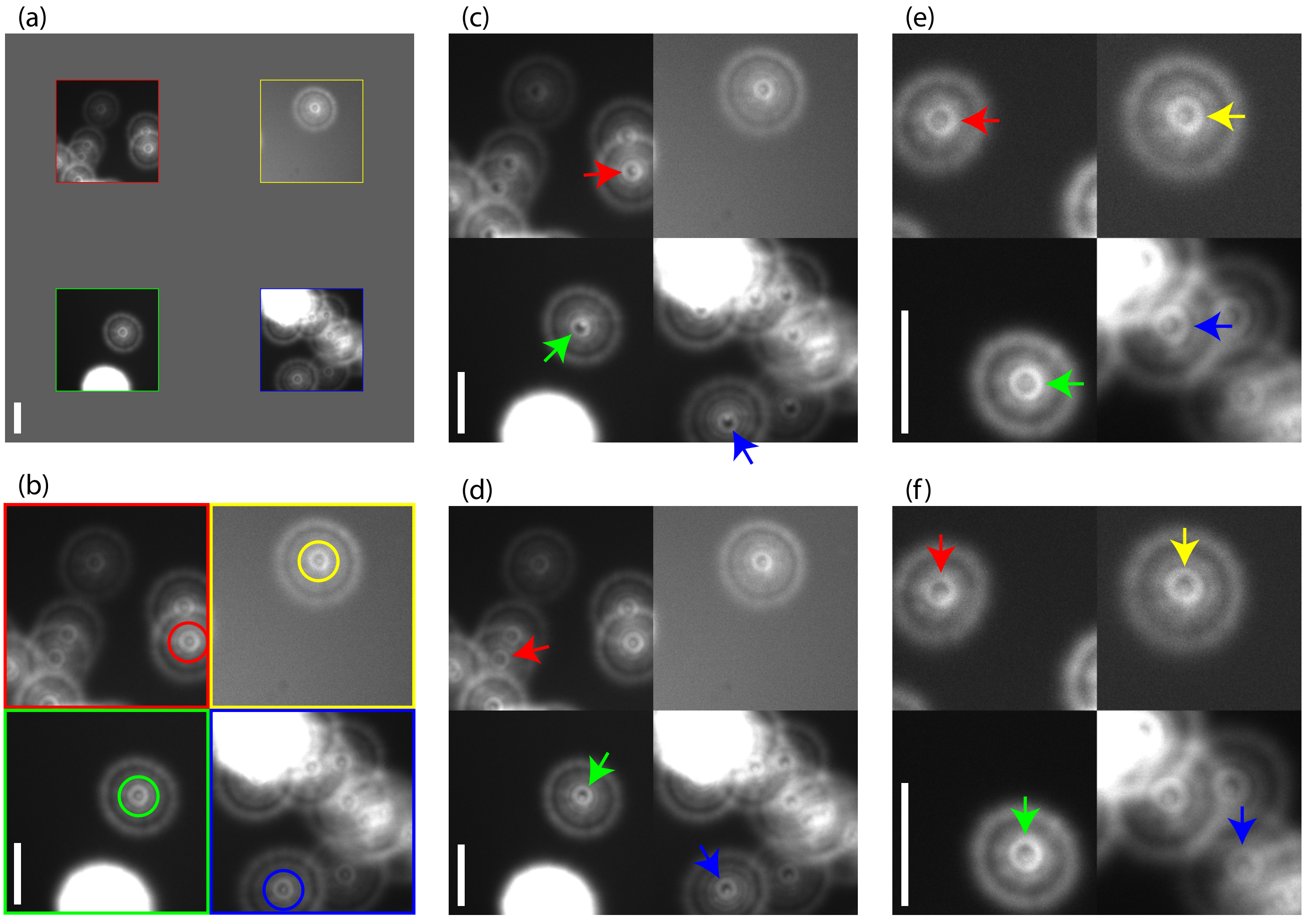


**Fig. S6.** Vortex phase plate alignment. (a) Selected imaged regions of defocused beads on the camera. (b) Zoom in on the four selected regions. The circles highlight the central region where a central peak is surrounded by a bright ring. The bright ring moves as the vortex phase plate is moved, in this image the center of the rings and the peaks coincide and thus the phase plate is properly aligned. (c) Vortex phase plate is too close to the microscope as the rings created by the vortex phase plate are too far radially outward. The arrows indicate the direction in which the rings should moved. (d) Vortex phase plate is too close to the camera when the rings are too far radially inward. (e) When the vortex phase plate is aligned along the optical axis all spots should look the same throughout the FOV. The position can then be fine-tuned by shifting horizontally. (f) Vortex phase plate slightly misaligned vertically. Scale bars are 10 $\mu$m.

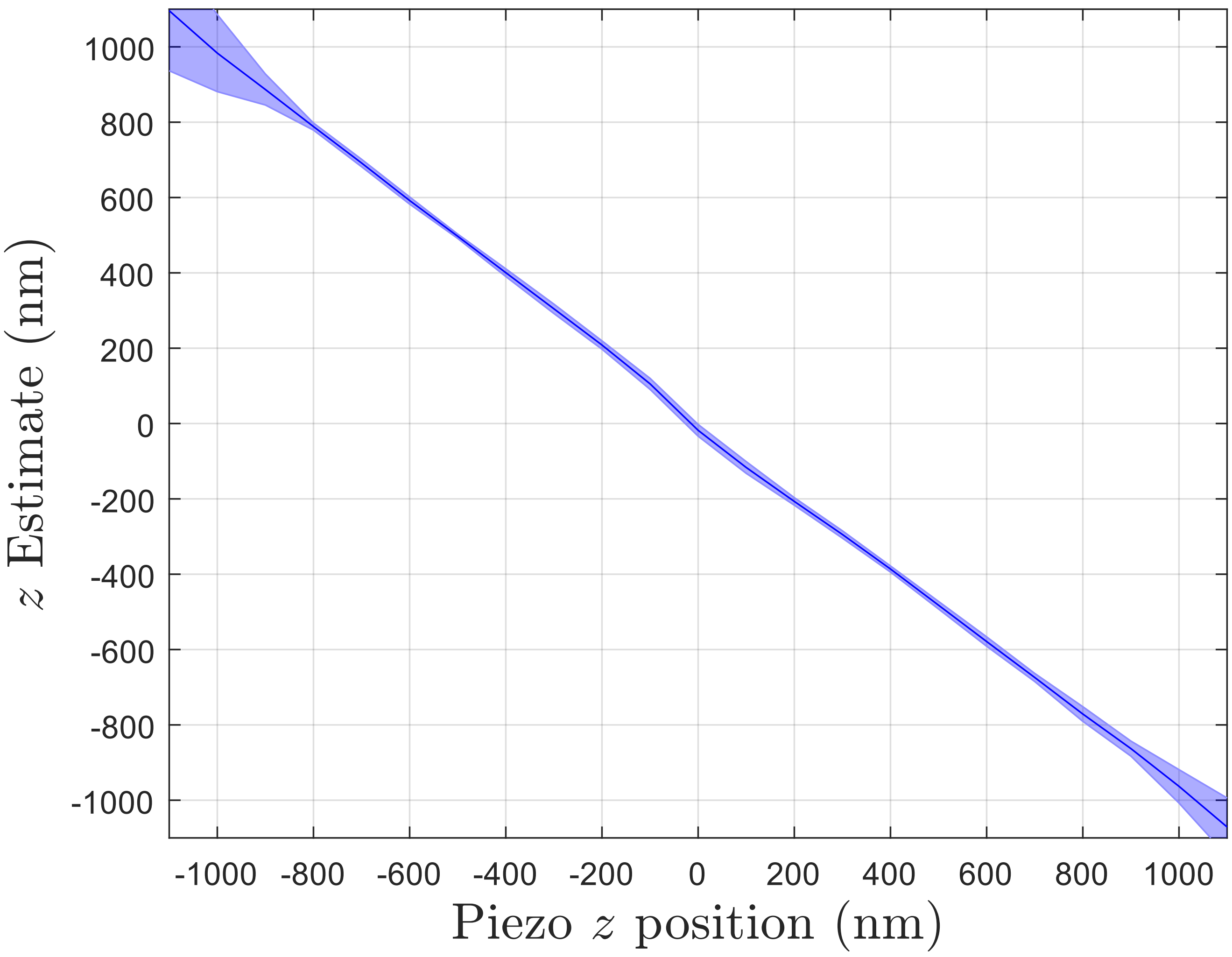


**Fig. S7.** Mean estimated $z$ position as a function of the piezo $z$ position with the shaded region representing $\pm\sigma$. The average is taken from 37 single molecules of varying orientations where the piezo $z$ is realigned in processing to account for the in focus position not corresponding exactly with $z_{\text{piezo}} = 0$. Each of the 37 molecules is fitted with a linear polynomial for $|z| \leq 600$, resulting in an average slope of $-0.99 \pm 0.01$ and a Root Mean Square Error (RMSE) of $16$ nm. The slope is negative due to the opposing definition of the piezo stage $z$ and the sample $z$.

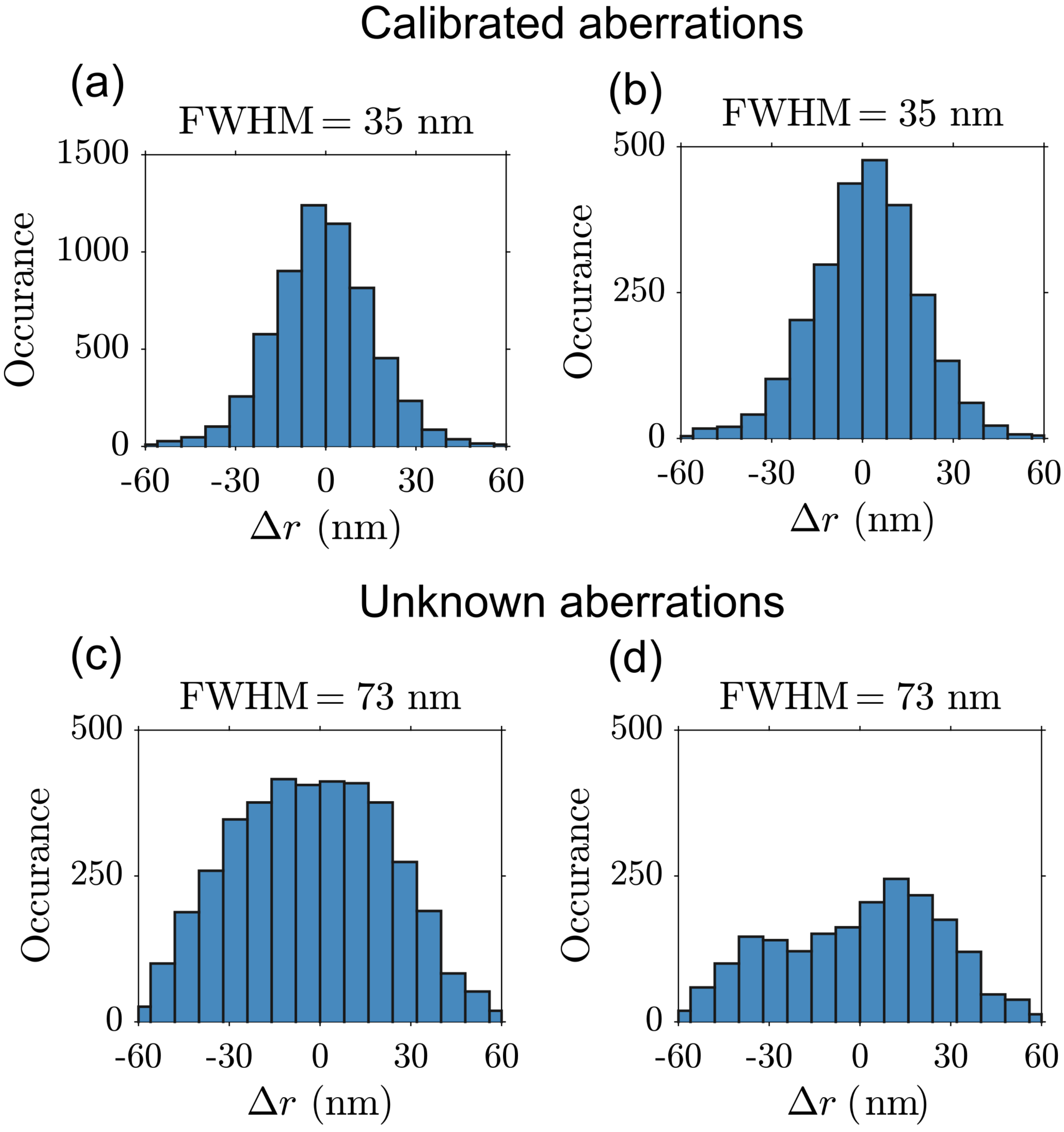


**Fig. S8.** Affect of calibrated PSF model for the localizations of the $\lambda-$DNA in the main text [Fig. 4(a)]. (a) Position deviation from the spline fit to the DNA axis using a calibrated PSF model results in a Gaussian-like distribution with $\mathrm{FWHM} = 35$ nm and (b) for polar angles $45 \leq \theta \leq 135$ results in a similar distribution with $\mathrm{FWHM}35$ nm. (c) Using a PSF model without calibrated aberrations results in a broad distribution with $\mathrm{FWHM} = 73$ nm and (d) for polar angles $45 \leq \theta \leq 135$ results in a non-uniform distribution with $\mathrm{FWHM} = 73$ nm. (a-b) The FWHM is evaluated using a normal distribution fit with support $\Delta r = \pm 40$ nm and (c-d) a non-parametric fit with a normal kernel.

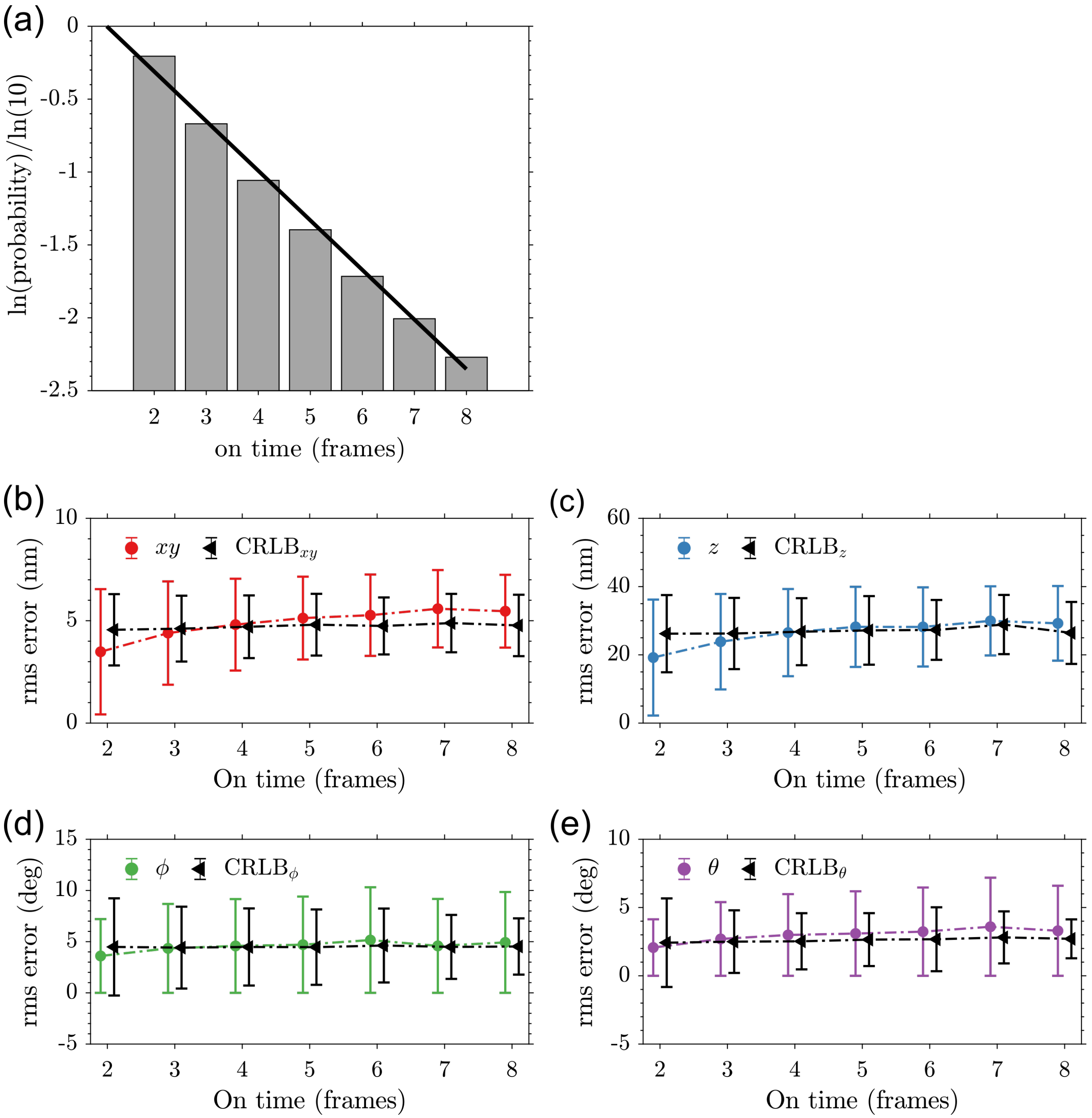


**Fig. S9.** Single-molecule run length analysis in $\lambda$-DNA experiment. (a) Natural-logarithm probability-distribution (gray bars) of subsequent on-time events up to 8 frames for the same emitter. The average on-time over all events is 1.5 frames, whereas the fitted exponential distribution gives an on-time of 1.3 frames (black line). (b) Lateral localization error (root-mean-square value) and CRLB (mean and s.d.) determined from repeated localizations. In the same way, (c) axial localization error, (d) azimuth angle error and (e) polar angle error estimated from repeated localizations. The estimated RMS error matches well with the estimated CRLB for all parameters and the number of on-time events. The total number of linked on-events is 302,541 with experimental conditions and data analysis as further specified in section 5 and 6, respectively.

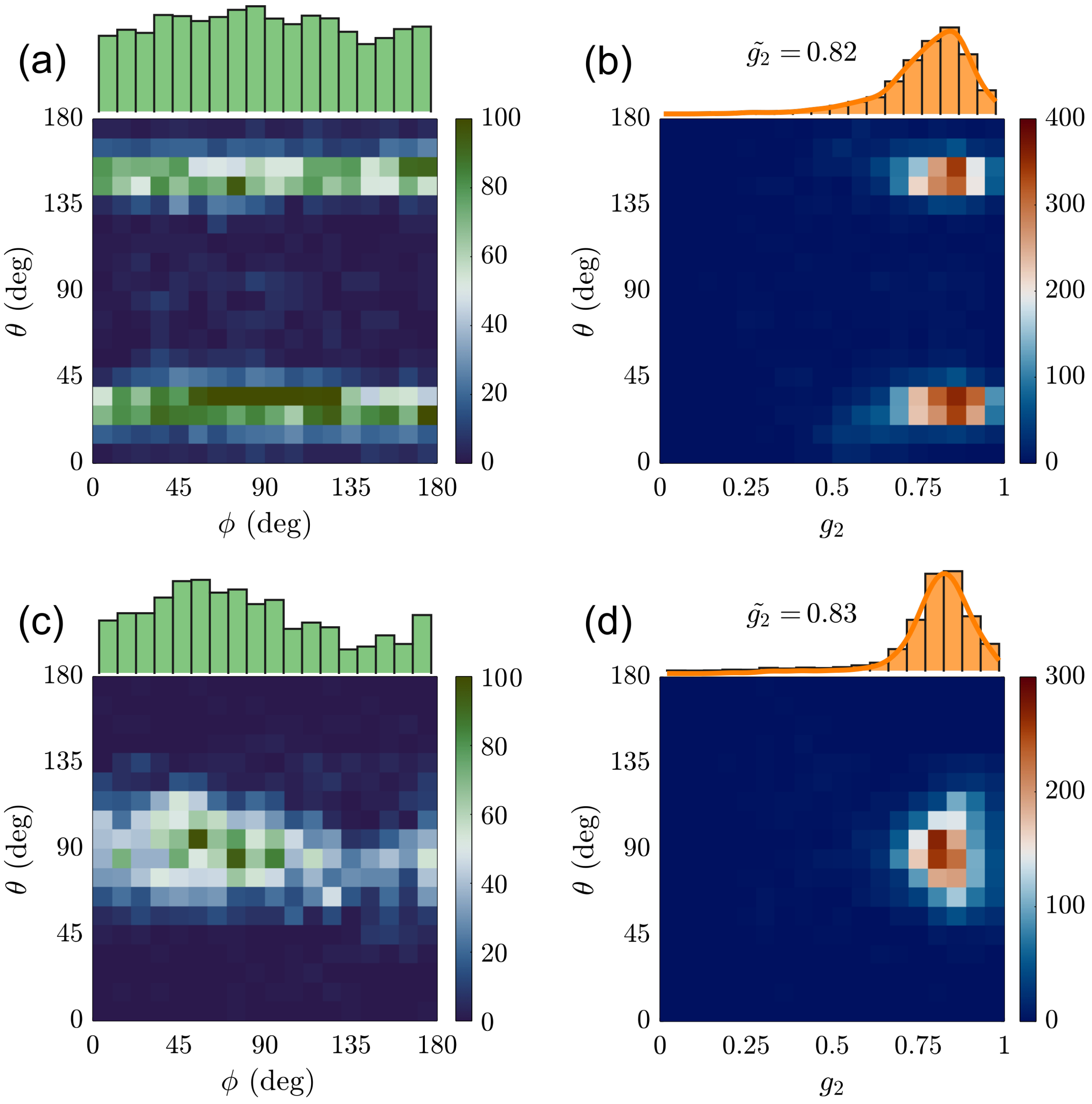


**Fig. S10.** Orientation and diffusion estimation on fixed molecules. (a) Bivariate histogram of the azimuth angle ($\phi$) versus polar angle ($\theta$) and (b) diffusion coefficient ($g_2$) versus polar angle as estimated on single-molecules under TIRF illumination directly spin-coated onto a coverslip. The marginal histograms show the azimuth and diffusion distribution, respectively, with the median diffusion coefficient specified in the plot. (c-d) The same as in (a) and (b), but estimated on single-molecules embedded in a thin layer of PMMA under epi-illumination . In both these experimental cases with different orientation distributions, there is no correlation between the estimated parameters. The number of single-molecules analyzed is 7042 in (a-b) and 4034 in (c-d) with experimental conditions as further specified in section 5.
